# Distinct age- and pathology-dependent epitranscriptome and translational dysfunction in a tauopathy mouse model

**DOI:** 10.1101/2025.05.15.654371

**Authors:** Guangxin Sun, WeiWei Lin, Ruixi Chen, Lulu Jiang, Michael J. DeMott, Abigail McShane, Chi Kong Chan, Dylan Ehrbar, Xiaoling Zhang, Thomas J. Begley, Andrew Emili, Benjamin Wolozin, Peter C. Dedon

## Abstract

Chronic neurodegenerative diseases, such as tauopathies, cause major metabolic changes in the brain affecting gene expression at both the transcriptional and translational levels. Our understanding of how regulation of translation changes with disease has focused on mRNA and its translational regulatory factors, RNA binding proteins, and microRNAs, despite clear evidence for translational and post-translational dysfunction in ADRD and tauopathies. The neurobiology of tRNA has only recently begun to be studied, but the impact of chronic neurodegenerative diseases on tRNA biology and translational dysfunction is largely unknown. We have previously shown that the tRNA pool and tRNA modifications behave as a system to regulate the cellular stress response by undergoing stress-specific reprogramming and causing selective translation of mRNAs from codon-biased stress response genes. Here we tested this stress-induced tRNA reprogramming and codon-biased translation system in the response to mutant tau expression by performing mass spectrometric quantification of ∼8500 proteins and 49 tRNA modifications, AQRNA-seq analysis of 222 cytosolic and mitochondrial tRNAs and other small RNAs, and informatic analysis of codon usage patterns in >23,000 protein-coding genes. Analysis of these datasets revealed that aging and tauopathy elicit major adaptation of the tRNA transcriptome and epitranscriptome as well as corresponding evidence of a program of translation of families of codon-biased genes for aging and disease responsive proteins. The mitochondrial tRNA transcriptome showed a strong response to aging and disease with 21 of the 22 mt-tRNAs showing age and disease-linked increases in expression, accompanied by mitochondria-specific modifications such as ms2i6A and f5C. Surprisingly, there were few significant changes in the 203 cytosolic tRNA isodecoders. However, a 10-fold increase tRNA isodecoder tRNA-Arg-TCT-5-1 was accompanied by increased translation of proteins encoded by genes highly enriched in its AGA cognate codon. These changes in tRNA biology are mirrored by strongly biased use of synonymous codons among the most highly upregulated and downregulated proteins in the P301S MAPT mice. Taken together these findings suggest the aging and disease brain produces an integrated response for translational control that is highly integrated with changes in tRNA biology.

## Introduction

Tauopathies are neurodegenerative diseases that are characterized by the accumulation of neurofibrillary tangles, which are composed of aggregated microtubule associated protein tau (Tau) and accumulate in neurons.^1^ The most common form of tauopathy is Alzheimer’s disease (AD), which is characterized by two pathological hallmarks, neurofibrillary tangles and β-amyloid plaques, which are composed of aggregated β-amyloid (Aβ) and accumulate in the extracellular milieu. Tauopathies, particularly AD, are heterogeneous and can exhibit accumulation of multiple other types of protein aggregates.^1^ The accumulation of neuropathology is associated with extensive neuroinflammation.^1, 2^ Together these factors produce chronic stress and cause the neurodegeneration that gives rise to cognitive loss.

Biochemical and proteomic studies show many changes occurring in the degenerating brain, including striking changes in levels of many proteins and strong changes in post-translation modifications, such as phosphorylation.^3, 4^ Proteins such as Clu, C1q and Gfap show strongly increased expression as the disease progresses. Other proteins, such as Tau, show dramatic changes in phosphorylation patters, some of which are thought to drive the disease process.^4, 5^ The brain responds to the stress of neurodegeneration with a strong translational stress response.^6^ The translational stress response is manifest by the accumulation of RNA membraneless granules, termed stress granules, and also by inhibition of protein synthesis.^6^ These studies of stress granules and RNA translation capture important aspects of the pathophysiology of neurodegeneration, but overlook other key elements of the stress response.

Increasing evidence indicates that the pool of transfer RNAs (tRNAs) and the 50+ modified ribonucleosides comprising the epitranscriptome represent another major aspect of the translational stress response. The universal genetic code provides 61 codons specifying 20 amino acids, which implies a redundancy of synonymous codons for many amino acids.^7^ However, this simple description does not capture the true complexity of tRNA pathways because, for every isoacceptor tRNA that reads one synonymous codon, there are up to 10 variant tRNA sequences, termed isodecoders, that read the same codon.^7^ With 500 total tRNA genes and about 250 expressed at any one time,^8^ the isodecoders vary in their expression among tissues and also vary in the changes in expression in response to stress.^9, 10^ Every tRNA isodecoder can also be modified by up to 50 post-transcriptional modifications.^11, 12^ We have shown that the tRNA pool and tRNA modifications behave as a system to regulate the cellular stress response by undergoing stress-specific reprogramming and causing selective translation of mRNAs from codon-biased stress response genes.^12–15^ How the tRNA system responds to the stress of chronic brain disease, though, is largely unknown.

There is emerging evidence for critical roles of the tRNA pool and tRNA epitranscriptome in brain function and disease.^16–18^ For example, Kapur et al. profiled tRNA expression in a dozen different neuronal cell types in mice, observing that neuron-specific homeostasis and protein translation depend on tRNA pool composition reflecting codon demand.^10^ Blaze et al. observed that loss of Nsun2, which inserts m5C in tRNA, mRNA and other RNAs, reduced tRNA m5C levels, reduced expression of tRNA-Gly isodecoders, decreased synaptic signaling at pyramidal neurons, and defective memory.^19^ With regard to Alzheimer’s disease and related disorders, Pereira recently reported that expression of ELP3, which carboxymethylates wobble uridines in tRNA to form cm5U, is reduced in brain tissue from ADRD patients and amyloid mouse models, which correlates with increased amyloid plaque density.^20^

Here we examine the role of tRNA reprogramming and codon-biased translation in the PS19 P301S Tau mouse model of tauopathy.^21, 22^ In this model, heterozygous P301S(+/−) mice exhibit 5-fold over-expression of a disease-linked mutant human tau isoform (4R P301S Tau protein), while wild-type (WT) P301S(−/−) mice lack the mutant tau gene. With increasing age, the mouse model develops increasing tau pathology, leading to the appearance of strong tau pathology by 6-9 months with concomitant synaptic neurodegeneration, progressing to cell death and then death of the mouse. We used this mouse model to characterize the role of the protein translation system in the progressive accumulation of neuropathology and progressive neurodegeneration. We now report that the neurodegenerative process produces striking age- and pathology-dependent changes in the types and amounts of RNA modifications at 9 months, accompanied by substantial ribosome degradation. While the levels of 203 expressed tRNA isodecoders remained relatively constant, we observed distinct codon biases among the pathology-dependent increases and decreases in protein levels. These results point to age- and pathology-dependent reprogramming of the tRNA pool by RNA modification writers and erasers, which shows strong links to changes in the translation of mRNAs selectively translated by the reprogrammed tRNA pool.

## Results

### Development of pathology in the PS19 P301S MAPT mouse line

The PS19 P301S MAPT mouse is a well-documented mouse line that develops tau pathology, astrogliosis, inflammation, and neurodegeneration.^21, 22^ P301S(+/−) mice exhibit about 5-fold over-expression of human P301S Tau (**Fig. 1A**). Pathological Tau becomes evident by immunostaining at 6 months with the appearance of phosphorylated Tau (pS202/205) and misfolded Tau (detected with the MC1 antibody) (**Fig. 1B, C**). The amount of pathology increases steadily being robust by 9 months of age (**Fig. 1B, C**). Astrogliosis detected by labeling of Gfap and inflammation detected by labeling with Iba1 both become increasingly prominent from 6 – 9 mo (**Fig. 1D**). Neurodegeneration occurs in parallel with the accumulation of neuropathology and, by 9 mo, can be readily detected by labeling of synaptophysin, MAP2, and cleaved Caspase 3 (**Fig. 1E**). Subsequent proteomics analyses of brain hemisections at 3, 6, and 9 mo used cohort sizes of 4 males (M) and 4 females (F) for wild-type (WT) P301S(−/−) mice and tau over-expressing (mutant) P301S(+/−) mice, while epitranscriptome and ribonucleome analyses in brain hemisections were performed with the following cohorts: WT at 3 mo, 7 M, 6 F; Mutant at 3 mo, 2 M, 10 F; WT at 6 mo, 2 M, 10 F; mutant at 6mo, 7 M, 6 F; WT at 9 mo, 2 F; mutant at 9 mo, 2 M (**Suppl. Table S1**).

### Proteomic analysis of the tauopathy mouse model reveals strong upregulation of inflammatory proteins

To test the role of translational regulation of gene expression in the PS19 mouse model of tauopathy, we performed a quantitative proteomics analysis with extracts from brain hemi-sections. **Suppl. Table S2** details the proteins with significant changes in the mutant P301S(+/−) mice compared to the WT P301S(−/−) mice.

An initial quality control assessment revealed an average of 8,579 proteins among each cohort. There was a robust distribution across Gene Ontology (GO) categories, including the expected ADRD/tauopathy gene families (**Suppl. Fig. S1**). Comparing mutant to WT mice, we observed an increasing number of up- and down-regulated proteins at p<0.1 as a function of disease progression from 3 mo to 9 mo (1222-1337 up, 1130-1567 down; **Suppl. Table S2**).

Gene Ontology (GO) category analysis of the most up-regulated proteins across the time course revealed functional changes strongly reflecting tauopathic mechanisms of cell and tissue damage (e.g., mitophagy, autophagy, neurodegeneration), the neuroinflammatory response to the damage (e.g., microglial activation; Parkinson, Huntington, ALS, prion diseases), and the activation of cell regeneration pathways reflecting the epithelial–mesenchymal transition (EMT; e.g., adherens and tight junction, focal adhesion, cytoskeleton) (**Suppl. Fig. S1**).^23–25^ Analysis of individual proteins reveals gene expression consistent with the pathology of mutant tau protein expression, including high levels of MAPT at all time points (**Suppl. Fig. S1; Suppl. Table S2**). Inflammatory genes stood out among the proteins most increased in the mutant mice compared to WT at 9 months, corresponding to 11 of the top 20 up-regulated proteins: C1qa/b/c, C4b, Ifit3, Ifitm3, Cd14, Lyz2, Lag3, and Bst2 at p<0.05; and Sec61g at p<0.1 (**Suppl. Fig. S1, Suppl. Table S2**). Interestingly, of these proteins, only C4b was detected among the 1232 and 1293 upregulated proteins at 3 mo (p<0.1) and 6 mo (p<0.05), respectively. This is consistent with the immunohistologic time course of neurodegeneration, which was detected at 9 mo (**Fig. 1E**).

The importance of neuronal damage and inflammation caused by over-expression of the mutant tau proteins was also apparent in the proteome and phospho-proteome. Notably, the transcription factor Zeb1, master modulator of neuroinflammation and the EMT response to neurodegeneration,^23^ was the most highly differentially phosphorylated protein in the PS19 mice other than MAPT; phosphorylation of Zeb1 correlates with activation of the inflammatory system.^23, 24^ Similarly, the dual function Trmt13^25^ was among the most highly up-regulated proteins at 3 and 6 months (**Suppl. Fig. S1, Suppl. Table S2**). Trmt13 is important because it plays a strong role in regulating tRNA metabolism, installing 2’-O-methyl groups at stem loop position 4 in the tRNA isodecoders for Gly-GCC, Pro-UGG, Gly-CCC, Pro-CGG, Gly-UCC, and Pro-AGG^25, 26^. At the same time, Trmt13 functions as a transcriptional co-activator of EMT in tissue repair as well as inflammation, which is critical to the pathophysiology of tauopathies^25, 27–29^.The increased activity of the Zeb1 and Trmt13 regulatory pathways reflects the full scope of cell and tissue damage, neuroinflammation, and cell regeneration pathways.^23–25^ For example, an important mediator of neurodegeneration that shows up in the earlier time points was the dual specificity mitogen-activated protein kinase kinase 5 (Map2k5; **Suppl. Fig. S1, Suppl. Table S2**), which serves as a scaffold for the MAP3K2/MAP3K3-MAP3K5-MAPK7 signaling complex.^30, 31^ Map2k5 contributes to the pathophysiology of Parkinson disease, being genetically linked to restless leg syndrome in PD^32^ and contributing to dopamine deficiency.^33^ Map2k5 also contributes to TDP-43-mediated toxicity by inhibiting autophagy.^34^ Oxytocin (Oxt) shows up as strongly increased at the early stages of the PS19 mouse model (**Suppl. Fig. S1, Suppl. Table S2**), which is consistent with a general neuronal stress response and activation of the hypothalamic-pituitary-adrenal (HPA) axis (HPA) in ADRD and tauopathy.^35, 36^ Other processes reflecting neurodegeneration were evident at the 3- and 6-month time points, including multiple members of the large solute carrier family (Slc)^37^ among the 20 most down-regulated proteins: the neuro-protectant Slc1a3 excitatory amino acid transporter 1 (EAAT1, glutamate), the Slc38a3 gene encoding the sodium-coupled glutamine acid transporter 3 (SNAT3), and the Slc8a3 dopamine transporter (**Suppl. Fig. S1, Suppl. Table S2**).

The proteomics data are also consistent with both age- and tauopathy-dependent mitochondrial pathology.^38, 39^ Mitophagy resolution of damaged mitochondria and compensatory mitobiogenesis in response to neuronal stress and tissue damage are well established pathways activated in aging and neurodegeneration.^38, 40^ The master transcriptional regulator of mitobiogenesis, Nrf1,^41^ was strongly up-regulated in tauopathy at 3 and 6 mo, becoming down-regulated by 9 mo (**Suppl. Table S2**). While highly down-regulated at 3 mo and 6 mo in mutant mice, the mitobiogenesis protein translocases Tomm40 and Timm50^42, 43^ and the brain-enriched mitophagy regulator Atg9a^44, 45^ were both among the 100 most up-regulated proteins at 9 mo (**Suppl. Table S2**). Increased Tomm40 expression is associated with decreases in mitochondrial function and DNA copy number.^46^ While aging is associated with a reduced mass of functional mitochondria, increased damaged mitochondria, and increased mitochondrial size in neurons,^47^ there is an increase the number of damaged and dysfunctional mitochondria as well as increase the diversity of mitochondrial fragments and large mitochondria in AD and neurodegeneration.^48^ These observations are consistent with the up-regulation of mitochondrial fission protein Fis1 and the fusion protein Mfn1in mutant compared to WT at 9 mo. Ribosomal proteins present another mitochondrial distinction between aging and neurodegeneration, with an increase in the density of nearly all cytosolic ribosomal proteins as function of age in both mutant and WT mice, while mitochondrial ribosomal proteins increased with age in WT mice and plateau or decrease at 9 mo in tau-expressing mice (**Suppl. Fig. S2**). This behavior matched the differential expression in the mutant versus WT mice, with mitochondrial ribosomal proteins Mrps17 and Mrps18c among the most highly up-regulated proteins at 3 and 6 mo in the mutant mice, becoming down-regulated at 9 mo (**Suppl. Table S2**). Overall, these observations are consistent with a complicated picture of mitochondrial damage, reduced numbers of functional mitochondria, and increases in mitochondrial components involved in mitobiogenesis. This plays out with mitochondrial tRNAs and the modifications as discussed shortly.

There are several striking tau-dependent shifts in neurofunctional proteins during the 3- to 9-month time course representing either neuroregeneration or elevated compensatory activities in existing cells. For example, 24-39 different solute carrier (Slc) proteins were detected at each of the three time points, with a dramatic shift from mostly under-expressed at 3 mo and to mostly over-expressed in the mutant mice at 9 mo (p<0.1; **Suppl. Fig. S2; Suppl. Table S2**). Even more striking was the Rab family of GTPases that regulate intracellular membrane trafficking, such as neuronal vesicle transport.^49^ Of the 28 total Rab proteins detectable in the proteomics analysis, 25 shifted from down-regulated at 3 and 6 mo in the mutant mice to up-regulated at 9 mo (p<0.1), with the other 3 proteins (Rab26, Rab11fip2, Rab34) showing the opposite behavior (p<0.1; **Suppl. Fig. S2; Suppl. Table S2**). Overall, the proteomics changes reflect those classically observed in human neurodegenerative diseases. The systematic and wholesale changes in functional gene families, while reflecting the physiological mechanisms underlying tauopathy, raised the question of the molecular mechanisms underlying the large shifts in gene expression, mechanisms linked to very strong biased use of synonymous codons in genes for proteins up-regulated in tauopathy.

### Proteomic analysis reveals tauopathy-dependent codon-biased translation

We proceeded to interrogate the proteomics data to determine whether the changes in protein levels were associated with codon-biased translation.^12, 14, 15, 50^ The first step in this process was to quantify codon usage patterns across the mouse genome using an isoacceptor-based codon counting algorithm^51^; the resulting data are presented in **Supplementary Table S3**. Partial least-squares regression analysis of codons enriched in genes for the 20 most up- and down-regulated proteins are shown in **Figure 2**. The three scores plots in **Figure 2** (panels A, C, and E) show clear discrimination of up- and down-regulated proteins at each age, with the loading plots (panels B, D, and F) detailing the codons enrichments associated with the differentially regulated proteins. For example, up-regulated proteins at 3-and 6-mo show robust enrichment in Leu-CTA, while the 20 most down-regulated proteins are enriched in the synonymous counterpart Leu-CTC (**Fig. 2A,C**). This behavior reverses at 9-mo, with up-regulated proteins in the mutant mice enriched for Leu-CTC, while the 20 most downregulated proteins are enriched for Leu-CTA (**Fig. E, F**). Similar differential enrichments of synonymous codon partners are observed for many other codons at each age (**Figure 2**).

**Figure 2.**
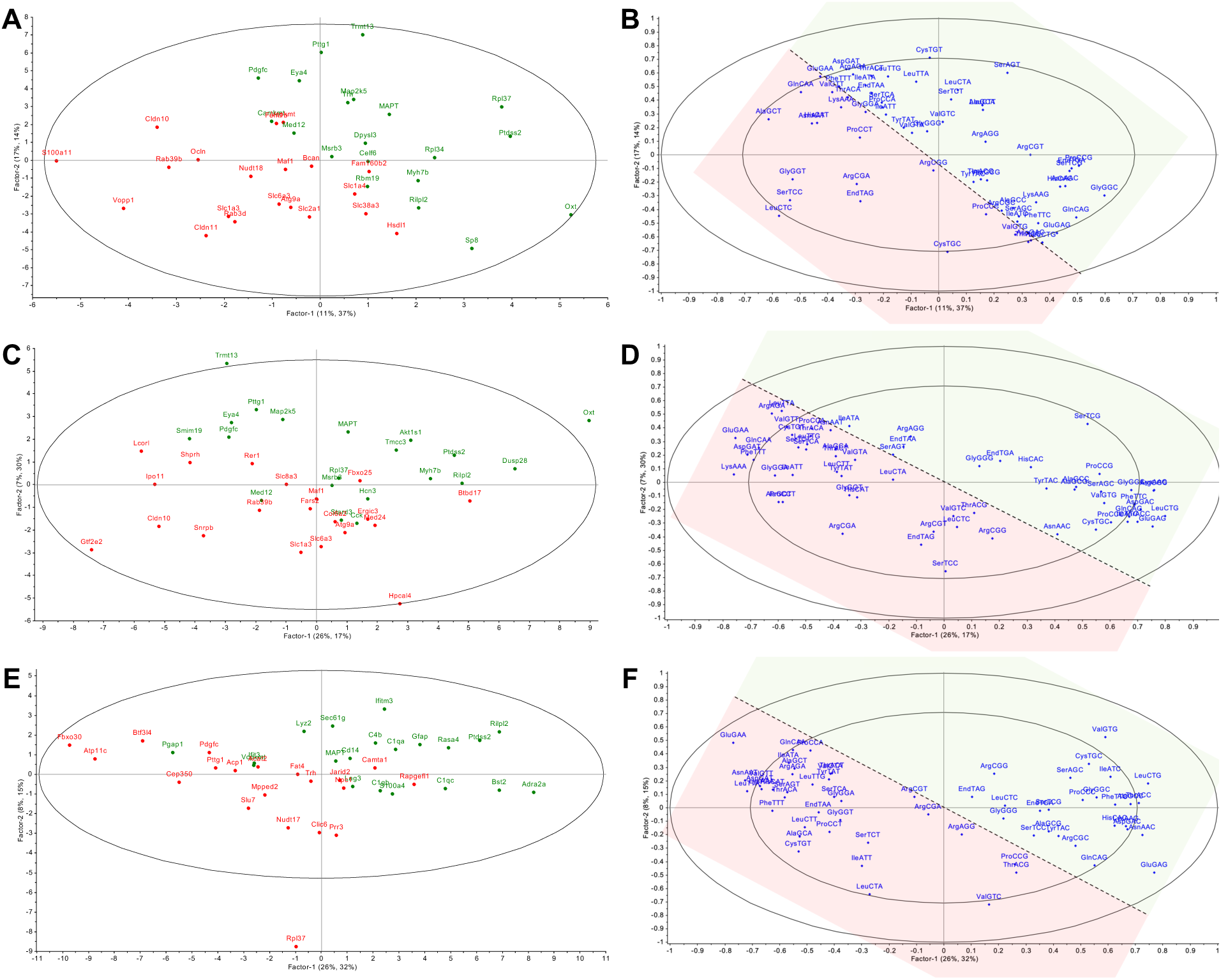
Expression of mutant tau protein causes codon-biased translation. The scores plots (**A**, **C**, **E**) and loadings plots (**B**, **D**, **F**) were derived from a partial least squares regression (PLSR) analysis of codon usage patterns in the 20 most over-expressed (green font, shading) and 20 most under-expressed (red font, shading) proteins in brain hemisections from P301S(+/−) mice expressing mutant tau protein compared to P301S(−/−) wild-type mice at 3 mo (**A**, **B**), 6 mo (**C**, **D**), and 9 mo (**E**, **F**) of age. Proteomics and codon usage data were taken from **Supplementary Tables S2** and **S3**, respectively. Up- and down-regulated proteins are clearly distinguished by patterns of synonymous codon usage, some of which reflect specific tRNA modifications (Fig. 4) as discussed in the text.

The codon-biased translation is strikingly illustrated for groups of proteins differentially regulated by mutant tau expression in **Figure 3**. The Slc, Rab, and Tmem gene families, which are largely up-regulated in the mutant mice, show significant enrichments of specific synonymous codons. This is most notable for the Slc and Tmem families, with over-use of many G- and C-ending codons (**Fig. 3, Suppl. Fig. S3**). On the other hand, the Rab family over-uses A- and T-ending codons (**Fig. 3, Suppl. Fig. S3**). This differential codon enrichment is illustrated by a highly significant negative correlation between codons GluGAG, IleATC, LeuCTG, LysAAG, and PheTTC in Slc genes and the ArgAGA codon in Rab genes (**Suppl. Fig. S4; Fig. 3**; Pearson’s p<0.05). The role of codon-biased translation in the stress response to mutant tau expression is further demonstrated considering regulatory proteins noted earlier. For example, mitobiogenesis factor Nrf1 and Map2k5 both share codon usage patterns similar to the Rab family (A/T-ending codons) while the mitobiogenesis translocases Tomm40 and Timm50, and the mitophagy regulator Atg9a are all similar to the Slc family (G/C-ending codons and AGA).

**Figure 3.**
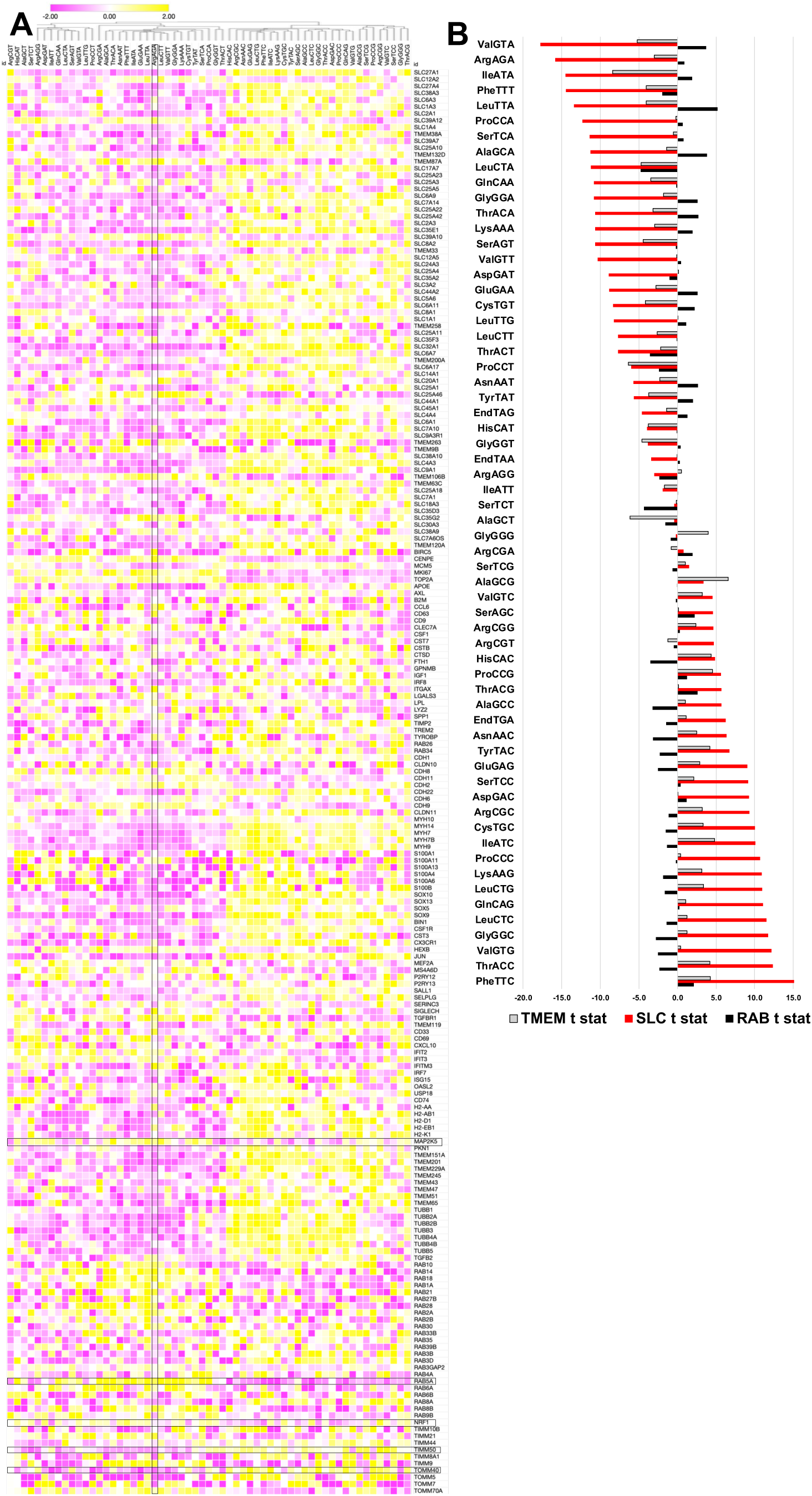
Codon usage patterns in genes up- and down-regulated by over-expression of mutant tau protein in the P301S mouse model. (**A**) Hierarchical clustering of Z-scores for codon usage (**Suppl. Table S3**) for examples of genes up- and down-regulated in mutant P301S(+/−) relative to wild-type P301S(−/−) mice. (**B**) T-statistics for codon usage in the Slc, Rab, and Tmem gene families differentially regulated by mutant tau protein over-expression.

This evidence for codon-biased translational responses to aging and tauopathy raised the question of links to changes in the tRNA pool, most notably for the tRNA-Arg-TCT isodecoders that read AGA.

### Disruption of the cytosolic and mitochondrial tRNA pools in tauopathy

To test the role of the pool of tRNA isoacceptors and isodecoders in the P301S tauopathy model, we quantified the levels for cytosolic tRNA isodecoders (cyto tRNA), mitochondrial tRNAs (mt-tRNAs), rRNA fragments, and miRNAs by applying AQRNA-seq to purified small RNA (<200 nt) extracted from mouse brain hemi-sections (**Suppl. Table S1**). AQRNA-seq provides a direct linear relationship between sequencing read count and molecular copy number by overcoming ligation biases and reverse transcriptase fall-off at RNA modifications.^52, 53^ The data processing summary and the raw and normalized read counts for the various RNAs are detailed in **Supplementary Table S4**. After trimming linkers, removing primer dimers, and deleting sequences shorter than 17 nt due to poor BLAST alignment, the “long read” portion of total reads represents biologically meaningful RNA molecules, with 75-87% assigned to tRNAs, rRNAs, and miRNAs, and the remaining 15-23% presumably representing unassigned RNA fragments from other sources (e.g., lncRNA) or fragments too small to assign (**Suppl. Table S4**).

AQRNA-seq analysis revealed 203 cyto tRNA isodecoders (47 isoacceptors) (**Suppl. Fig. S5**) and 22 mt-tRNAs (**Fig. 4C**), which is similar to the 224 tRNAs observed by Yu et al. in C57BL/6NCrl mice.^54^ The read counts, which are linearly proportional to copy numbers,^53^ ranged from 1 to 44,089, with read counts less than 10-20 viewed as too close to the detection limit to be statistically reliable (**Suppl. Table S4**). While the small cohort size at 9 months limits the statistical power of changes in RNA and modification levels, there are striking changes related to both age and tau pathology.

**Figure 4.**
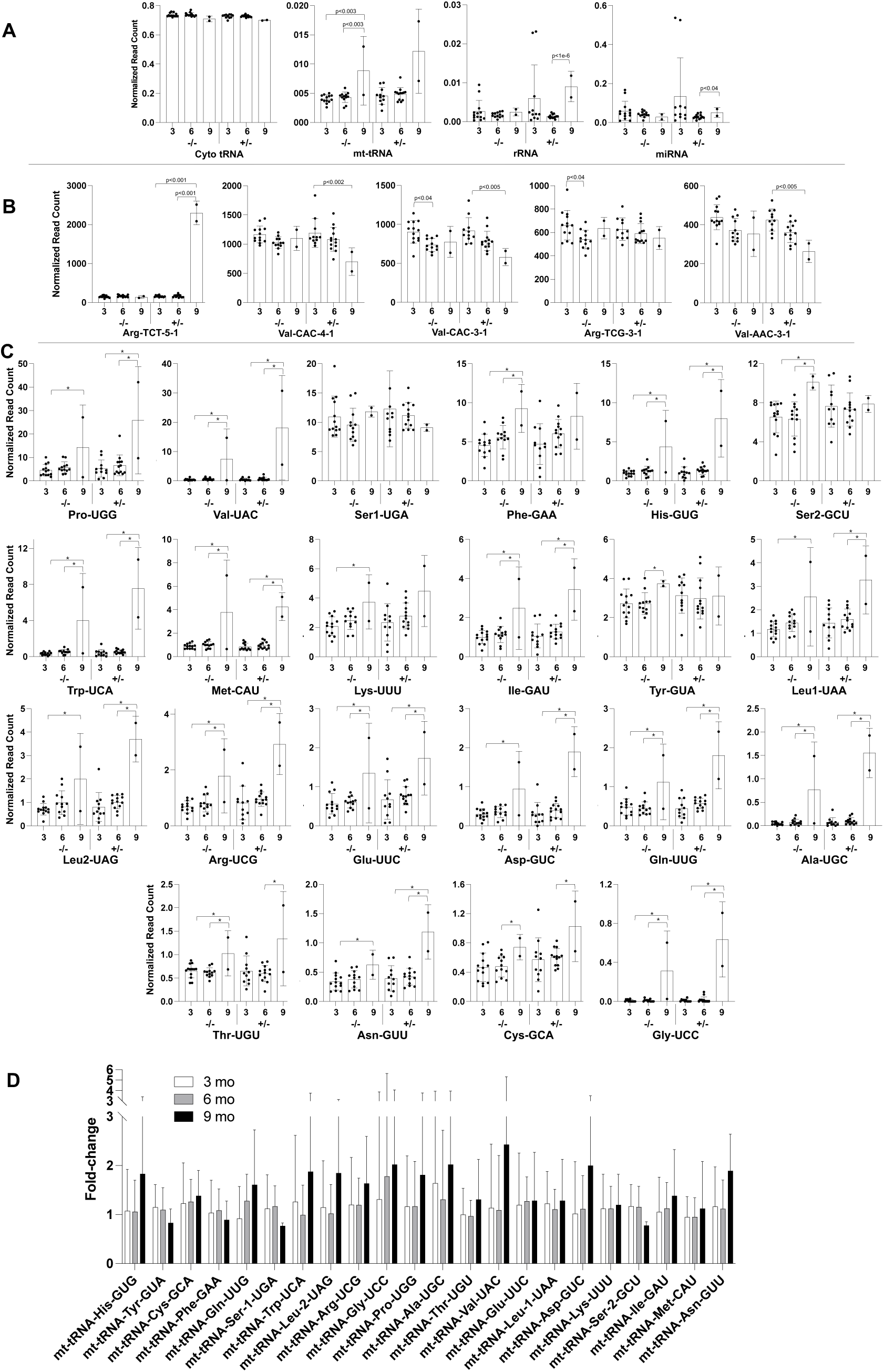
AQRNA-seq analysis of small RNA in mice expressin mutant tau protein. Small RNA extracted from mouse brain hemisections from P301S(−/−) WT mice and P301S(+/−) mice expressing mutant tau protein was subjected to AQRNA-seq to quantify tRNAs, rRNAs, and miRNAs. Analysis revealed 203 cytoplasmic tRNAs and 22 mt-tRNAs. (**A**) The pool of cytoplasmic tRNA isodecoders comprises 70-74% of assigned RNA species and shows little change as a function of age or pathology in the P301S mouse brain. Other RNA classes represented in the small RNA include mt-tRNAs (0.4-1%), 0.1-0.9% rRNAs (5S, 5.8S; fragments of 18S and 28S), and 3-13% miRNAs. (**B**) Among the few tRNA isodecoders changing significantly (asterisks denote t-test p<0.05) as a function of age and pathology is tRNA-Arg-TCT-5-1 that reads the AGA codon enriched in Rab genes. (**C**) While cytoplasmic tRNAs were largely unchanged, the mitochondrial pool (mt-tRNA) changes significantly (asterisks denote t-test p<0.05). (**D**) Fold-change values for mutant vs WT, while not statistically significant, showed upward and downward trends consistent with changes in the levels of their tRNA modifications. Data from **Supplementary Table S4**.

The analysis of tRNA levels revealed several novel behaviors, the first being a broad lack of significant changes in the levels of the 203 cytoplasmic tRNA isodecoders as a function of both age and pathology (**Fig. 4A, Suppl. Fig. S5**; **Suppl. Table S4**). The largest proportion of tRNAs showing >2-fold changes were all statistically insignificant and involved read counts <20 that likely represent noise due to the wide variance of the read counts. Even among the statistically significant changes (**Fig. 4B**), most of the changes are small (<50%). The exception to the generally small changes in the tRNA pool involves the large and selective increase in isodecoder tRNA-Arg-TCT-5-1 in the 9 mo P301S(+/−) mice, which was increased over 14-fold compared to all the other cohorts. Isodecoder tRNA-Arg-TCT-5-1 is one of 5 Arg-TCT isodecoders, with none of the other 4 tRNA-Arg isodecoders showing any changes associated with age or genotype (**Fig. 4B, Suppl. Fig. S5, Suppl. Table S4,**). Most mice express tRNA-Arg-TCT-4-1 as their main Arg-TCT isodecoder and this isodecoder is normally the most highly expressed in neurons.^10^ However, the background species for the P301S MAPT line, C57BL/6 mice, carries a mutation reducing levels of tRNA-Arg-TCT-4-1.^10^ The striking increase in isodecoder tRNA-Arg-TCT-5-1 might thus reflect a compensatory response to the dysfunctional isodecoder tRNA-Arg-TCT-4-1. It is important to note that the 9-mo cohort contained 2 male mutant P301S(+/−) and 2 female WT P301S(−/−) mice (**Suppl. Table S1**). Differential expression of the isodecoder tRNA-Arg-TCT-5-1 based only on sex could be ruled out because the 9 mo P301S(+/−) mice (male) showed elevated isodecoder tRNA-Arg-TCT-5-1 relative to the male P301S(+/−) mice at 3 and 6 months.

Another important feature in the AQRNA-seq data involves the mt-tRNAs (**Fig. 4C**). Compared to the relatively muted behavior of the cytosolic isodecoders, we found large statistically significant changes in most mt-tRNAs (**Suppl. Table S4**). The number and magnitude of the changes increased as a function of both age and pathology, with the largest changes occurring in the mutant P301S(+/−) mice at 9 mo, with 8 increasing between 2- and 13-fold and 6 decreasing by 40-70% at p<0.05 compared to 6 mo (**Fig. 4C**, **Suppl. Table S4**). There are no significant changes in the total number of normalized mt-tRNA reads, though there is a trend toward a decrease as a function of both age or pathology (**Fig. 4D**). While the changes in the mutant relative to the WT mice are not significant at p<0.1, there are clear trends both upward and downward in levels of mt-tRNAs as a function of both age and tau-induced pathology (**Fig. 4C**). It is important to point out that the mt-tRNAs comprise a total of 2200-3000 normalized reads compared to the 40,000-41,000 normalized reads for the cytosolic tRNAs (**Suppl. Table S4**), which implies that any associated mt-tRNA modifications will be diluted by the bulk of the cytosolic modifications.

AQRNA-seq also detects fragments of large RNAs resulting from degradation processes in the cells and tissues and, potentially to a small extent, from mechanical shearing during RNA isolation. As shown in **Figure 4A** and **Suppl. Table S2**, 5S, 5.8S, 18S, and 28S rRNA fragments were all detected in the small RNA fraction (<200 nt). Note that the 120 nt 5S rRNA variably partitions between large and small RNA fractions during purification and its behavior may not be biologically relevant.^55^ rRNA in the WT mice showed no age-dependent changes, ranging from 0.2-0.3% of total RNA reads, while there was a significant increase from 6 to 9 mo in the mutant mice (**Fig. 4A**); high variance in the 3 mo mutant mouse sample precluded meaningful comparison. We do not know the basis for the larger proportion of 28S rRNA fragments than 18S rRNA in all samples, but this is commonly observed and may be due to the nuclease-sensitive “hidden break” site in mammalian 28S rRNA.^56^ Regardless of the increase in rRNA in the 9 mo mutant mice, rRNA fragments accounted for at most 0.9% of total RNA <200 nt, compared to cyto tRNA at >70% (**Suppl. Table S4**), which has implications for interpreting age- and pathology-dependent changes in tRNA in modifications.

### tRNA and rRNA modifications change as a function of age and pathology

The results from analysis of translational codon bias (**Figs. 2,3**) and RNA quantification (**Fig. 4**) raised the question about changes in the RNA modifications of the epitranscriptome in the tauopathy mouse model. Here we quantified 49 modified ribonucleosides by LC-MS/MS in the same samples of small RNA extracted from brain hemi-sections used for AQRNA-seq analysis (**Suppl. Table S5**). The small RNA population comprises >70% tRNAs, along with miRNAs and rRNA and mRNA fragments (**Fig. 4A**).^53, 57, 58^ The resulting changes in the levels of the modifications as a function of both age and pathology are shown in the hierarchical clustering analysis and individual bar graphs in **Figure 5**. Despite the availability of only 2 brain samples for 9 mo mice, there are clear trends and large changes for many modifications, changes that are consistent with other datasets.

**Figure 5.**
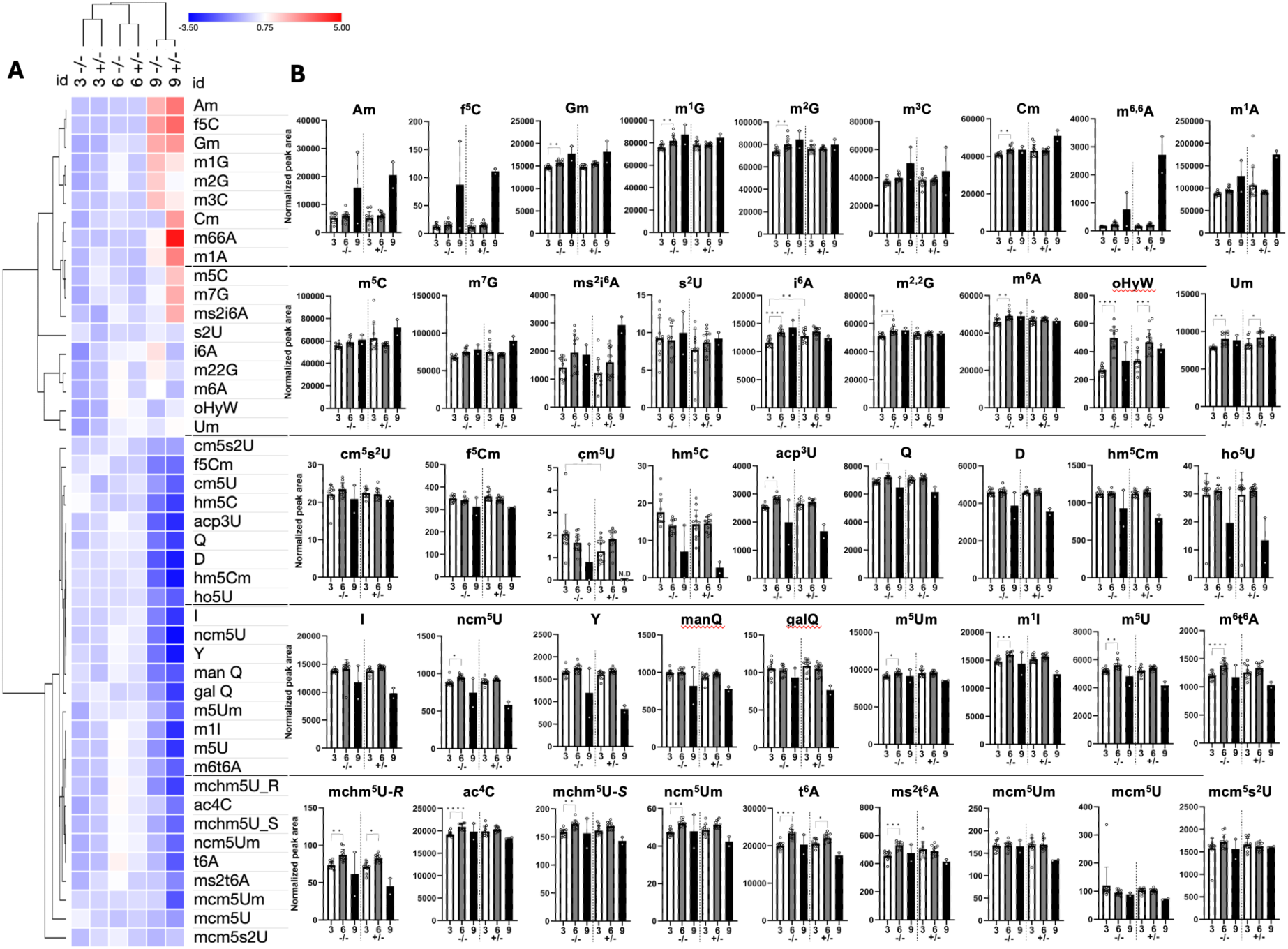
Changes in abundance of modified ribonucleosides in small RNA as a function of age and pathology in P301S tau-expressing mice. Small RNA (<200 nt) was from brain hemi-sections from P301S(+/−) mutant mice and P301S(−/−) wild-type mice and quantities of 45 modified ribonucleosides by LC-MS/MS. (**A**) Hierarchical clustering analysis age- and pathology-dependent changes in RNA modification levels. Color code reflects degree of change relative to a row average. The data reveal modest changes at 3 mo and 6 mo, with major age- and tauopathy-related changes in the levels of modifications occurring at 9 mo. (**B**) Graphs of normalized MS signal intensities for individual ribonucleosides. Data represent mean ± SD for 3 mo (N=12) and 6 mo (N=13) and deviation about the mean for 9 mo (N=2). Asterisks denote significance by Student’s T-test. Data from **Supplementary Table S5**

A general view of the heat map showed multiple significant changes in the levels of the modified ribonucleosides at all ages. The largest changes in RNA modifications were apparent at 9 mo, which roughly matches the time course for the severity of the pathology observed in the mouse brain.^21, 22^ Looking at individual ribonucleosides, the levels of Am, Gm, Cm, m66A, m1A, m5C, m7G, f5C, and ms2i6A increase at 9 months and, in some cases, increase in the P301S(+/−) mice expressing mutant tau protein more than in the WT P301S(−/−) mice (**Fig. 5**). Except for f5C and ms2i6A, the other modifications occur in tRNA, rRNA, and mRNA,^59–68^ which confounds assigning the source of these multi-functional modifications. For example, the increases in the rRNA and tRNA modifications Am>Gm∼Cm>Um from 3 to 9 mo in both WT and mutant mice (**Fig. 2**) are unlikely to be driven by the presence of rRNA fragments. The assignable rRNA fragments only account for 0.31% ±0.29% of the total RNA reads across all samples, with tRNA comprising 72±2%, mt-tRNA at 0.65±0.33%, and miRNA at 5.8±4.0% (**Suppl. Table S4**). Further, with Am, Gm, Cm, and Um as the most abundant modifications in tRNA, there are 41 Nm sites in 18S rRNA, 67 in 28S rRNA, and 2 in 5.8S rRNA^69^ for a total of 110 Nm in all rRNAs. This is similar to the ∼150 Nm in all human tRNA isodecoders^70^. Considering that rRNA and tRNA comprise 0.2-0.9% and >70% of all RNA species in the small RNA samples, rRNA only contributes only ∼1% of the Nm to the small RNA fraction. However, modification-containing rRNA and tRNA fragments could be present in the unassignable 15-30% of the “long reads”, which confounds the interpretation of the RNA degradation thus does not account for the increases in 2’-O-methylation modifications unless there is an abundance of unassignable degraded fragments containing modifications. The interpretation of the source of rRNA degradation products is complicated by two observations. First, the known translational shutdown caused by mutant tau expression^22, 71–73^ would be assumed to result in more ribosome degradation in mutant, which is consistent with the large decreases in ribosomal proteins detected at 9 mo in the mutant mice compared to WT mice (**Suppl. Table S2**). However, levels of rRNA fragments are similar in both WT and mutant mice (**Fig. 4, Suppl. Table S4**). Second, ribosomal proteins increased as a function of age in WT mice and increased in mutant compared to WT mice at 3 and 6 mo (**Suppl. Table S2**), which is consistent with an increase in ribosomes and rRNA. It is not known if an increase in steady-state levels of ribosomes would result in increased levels of rRNA fragments.

A similar argument applies for tRNA as the source of m1A increases at 9 mo in both WT and mutant mice (**Fig. 5**). m1A is among the most complicated RNA modifications, with 6 writers and 4 erasers targeting cytosolic and mitochondrial tRNAs and rRNAs (**Suppl. Table S6**), with TRMT6/TRMT61A for cytosolic tRNAs and TRMT61B and TRMT10C for mt-tRNAs.^74^ That rRNA fragmentation is not the source for increased m1A is supported by several observations: (1) there are only 2 sites for m1A in rRNAs (0 in 5S and 5.8S; 1 in 18S; 1 in 28S^75^) and (2) nearly all mammalian tRNAs are modified with m1A at position 58. While m1A inserted by mt-tRNA writer Trmt10C accounts for a small portion of total tRNA m1A^74^, Trmt10C is upregulated at 9 mo in mutant relative to WT mice (**Suppl. Table S2**). As discussed shortly, increased mt-tRNA containing m1A parallels increases in ms2i6A and f5C, which is consistent with the mitochondrial loss and bioenergetic stress^39^ involved in neurodegenerative diseases, which we propose triggers compensatory increases in mitochondrial tRNA modifications to support translation of core mitochondrial proteins, delay collapse, and maintain cell viability.

Among the modifications that decreased or trended downward as a function of age and tau pathology at 9 mo were the ELP-dependent wobble U modifications cm5U, cm5s2U, mcm5U, mcm5Um, ncm5U, and ncm5Um (**Fig. 5**); mcm5s2U was unchanged. This family of modifications arises from a complex series of reactions catalyzed variously by ELP3-5, CUT1/2, FTSJ1, and ALKBH8 as detailed in **Supplementary Figure S6**. These results parallel those of Pereira et al., who observed that ELP3 and its dependent wobble U modifications were (1) reduced in brain tissue from AD patients and an amyloid mouse model, (2) inversely correlated with the extent of amyloid plaque formation, and (3) positively correlated with proteotoxic stress.^20^ While the present studies show generally lower ELP pathway-dependent modifications, the levels of their catalytic enzymes did not show parallel changes, with variable up- and down-regulation as a function of age and pathology (**Suppl. Table S2**). This raises the potential for protein secondary modifications to regulate the activity of RNA-modifying enzymes.

For many tRNA modifications, age- and pathology-dependent changes in their levels are inconsistent with changes in the levels of their synthetic proteins. For example, t6A is synthesized by the KEOPS complex: Osgep (tRNA N6-adenosine threonylcarbamoyltransferase protein), Tp53rk, Tprkb, and Lage3/Pcc1. Keops uses a cofactor synthesized by Yrdc/Sua5. Both Yrdc/Sua5 and Lage3/Pcc1 increased at 6 and 9 mo in mutant compared to WT (**Suppl. Table S2**). The downward trend in t6A at 9 mo in the mutant occurs despite increases in Yrdc/Sua5 Lage3/Pcc1 in the mutant mice. These discrepancies between enzyme and modification levels suggest either that (1) there are other mechanisms for regulating enzyme activity, such as control of the activity of the synthetic enzymes by posttranslational modifications, or (2) the confounding effect of analyzing multiple tissue and cell types in the brain hemispheres analyzed here. However, there is a clear correlation between the changes in t6A level in WT mice (increase at 6 mo, decrease at 9 mo; **Fig. 5**) and the levels of the most abundant t6A-bearing tRNAs:^76^ tRNA-Arg-CCT-3-1/4-1, -TCT-1-1/2-1/5-1; tRNA-Lys-TTT-1-1/2-1; tRNA-Ser-GCT-2-1; tRNA-Thr-AGT-2-1/3-1/4-1, -CGT-1-1/2-1, -TGT-2-1 (Suppl. Fig. S5).

Further emphasizing the known importance of mitochondrial pathology and our finding of significantly altered mt-tRNA levels, we observed that 9 mo P301S(+/−) mice showed strongly increased levels of mt-RNA-specific ms2i6A, which is a member of the position 37 i6A family of modifications that also includes io6A and ms2io6A.^77^ ms2i6A is found on the mt-tRNAs Tyr-GUA, Ser-UGA, Phe-GAA, and Trp-UCA.^78^ This parallels the increase in ms2i6A-containing mt-tRNAs in both WT and mutant mice (**Fig. 4**).Perhaps the most striking change in tRNA modification levels involved hm5C and f5C, which are derived from m5C by progressive oxidation steps. m5C is present at 15 different locations on cytoplasmic tRNAs, inserted by NSUN2 at wobble position 34 and Dnmt2 at position 38,^66, 79, 80^ and at position 34 in mt-tRNA-Met by NSUN3.^81–83^ The fact that there are multiple enzymes catalyzing m5C formation at multiple sites in multiple cytoplasmic and mitochondrial tRNAs explains why m5C levels varied little as a function of age- and pathology in both mutant and WT mice (**Fig. 5**). It is at the stage of m5C oxidation to hm5C and f5C that distinctions begin to emerge for tRNA reprogramming and codon-biased translation in mouse brain. In mitochondria, Alkbh1 oxidizes m5C to hm5C and then to f5C in mt-tRNA-Met-CAU.^84^ Not found in cytoplasmic tRNAs, f5C34 enables mt-tRNA-Met-CAU to read both AUA and AUG codons;^85^ mt-tRNA-Met with m5C34 cannot decode AUA.^84^ This is important for mitochondrial translation since AUA encodes Met instead of Ile in mitochondria and mt-tRNA-Met is the only tRNA available for both translation initiation and elongation.^86^ Our observation of increases f5C in in both WT and mutant mice at 9 mo parallels the sharply increased levels of mt-tRNA-Met at 9 mo (**Fig. 4C**).

For cytosolic tRNAs, the story moves to 2’-*O*-methylated versions of hm5C and f5C, hm5Cm and f5Cm, respectively, and how their down-regulation in the P301(+/−) mutant may drive with shifts in codon-biased translation. FTSJ1 methyltransferase is responsible for formation of both f5Cm and hm5Cm,^84^ but was not detected in our proteomics analyses (**Suppl. Table S2**). f5Cm is located at the wobble position of tRNA-Leu-CAA^87^ and has been proposed to allow cytoplasmic tRNA-Leu-CAA to read both UUA and UUG codons for Leu.^84^ This is analogous to f5C allowing mt-tRNA-Met to read both AUA and AUG codons.^85^ Less is known about the location of hm5Cm except that it is (1) limited to small RNAs (<200 nt)^88^ and (2) highly abundant in mammalian cells at 1 per 5000 m5C.^89^ Suzuki and coworkers observed a broad distribution of wobble C modifications in the cytoplasmic isoacceptor Leu-CAA population (i.e., pooled isodecoders) from HEK293T cells: 5.7% f5C, 30% f5Cm, 34% hm5C, 26% hm5Cm, 2.6% m5C, and 1.9% C. We observed reductions of 15-80% in the levels of f5Cm, hm5C, and hm5Cm at 9 mo (**Fig. 5**) and insignificant changes in the levels of the four tRNA-Lue-CAA isodecoders (**Suppl. Fig. S5**), which suggests a role for Alkbh8 and other erasers in regulating tRNA modification levels.

## Discussion

Chronic neurodegenerative diseases, such as tauopathies, cause major metabolic changes in the brain affecting gene expression at both the transcriptional and translational levels. Our understanding of how regulation of translation changes with disease has overwhelmingly focused on mRNA and its translational regulatory factors, RNA binding proteins, and microRNAs, despite clear evidence for translational and post-translational dysfunction in ADRD and tauopathies.^1^ The neurobiology of tRNA has only recently begun to be studied,^10^ but the impact of chronic neurodegenerative diseases on tRNA biology and translational dysfunction is largely unknown. We have previously shown that the tRNA pool and tRNA modifications behave as a system to regulate the cellular stress response by undergoing stress-specific reprogramming and causing selective translation of mRNAs from codon-biased stress response genes.^12–15^ Here we tested this model in the P301S MAPT mouse model of tauopathy^21, 90^ by performing a multi-omic analysis of age-and disease-dependent changes in the tRNA transcriptome and epitranscriptome and how codon usage in mRNAs contributes coordinates with the tRNA changes to modulate protein synthesis in disease.

To test stress-induced tRNA reprogramming and codon-biased translation in the response to mutant tau expression, we performed mass spectrometric quantification of ∼8500 proteins and 49 RNA modifications in tRNAs, AQRNA-seq analysis of 222 cytosolic and mitochondrial tRNAs and other small RNAs, and informatic analysis of codon usage patterns in >23,000 protein-coding genes.^12–15^ Analysis of these datasets revealed that aging and tauopathy elicit major adaptation of the tRNA transcriptome and epitranscriptome as well as corresponding evidence of a program of translation of families of codon-biased genes for aging and disease responsive proteins. The mitochondrial tRNA transcriptome showed a broad dynamic response to aging and disease with almost half of the 22 mt-tRNAs showing age and disease-linked increases in expression. The changes in the tRNA epitranscriptome are similarly striking. Dozens of chemical modifications are observed affecting both cytosolic and mitochondrial tRNAs. Many of these modifications are known to affect tRNA biology, some of which change the codon specificity for the tRNA isodecoder. These changes in tRNA biology are mirrored by strong evidence of codon bias among the most highly upregulated and downregulated proteins in the P301S MAPT mice. Taken together these findings suggest the aging and disease brain produces an integrated response for translational control that is highly integrated with changes in tRNA biology.

One of the most striking features of these analyses is the strongly biased use of specific synonymous codons in genes encoding proteins highly upregulated in the stress of tauopathy (**Fig. 3**). The relationship of codon usage to protein synthesis focuses on a complementary arm of translational biology and identifies striking changes with disease. At each age, the 20 most upregulated and downregulated proteins exhibit use of synonymous, complementary codons. This codon bias is observed clearly when examining the SLC transporter family, which was one of the protein families exhibiting the strongest disease-linked codon bias (**Fig. 3**). GO ontology analysis shows that the members of this group exhibiting this bias are strongly linked to the endoplasmic reticulum, which well-known to play a critical role in the pathophysiology of tauopathies.^2, 91, 92^

The mitochondrial tRNA transcriptome and epitranscriptome stand out for their integrated and dynamic responses to aging and even more to disease. We observe an increase in multiple tRNA isodecoders that is evident in 9m WT mice and greater in 9m P301S MAPT mice. The increase in mt-tRNA isodecoders is matched by a correlated increase in mt-tRNA-specific modifications. For example, ms2i6A at position 37 of mt-tRNAs Tyr-GUA, Phe-GAA, Cys-GCA, Trp-UCA, and Ser-UGA facilitates base pairing and fidelity with A-ending codons.^77^ Several ms2i6A-containing mt-tRNAs were among the most highly upregulated at 9 mo (**Fig. 4**). These changes may be occurring to drive translation of mitochondrial genes, which have highly distinct codon usage patterns enriched for A-ending codons: Ser-UCA, Val-GUA, Glu-GAA, Leu CTA, Lys-AAA, and Leu-UUA (**Suppl. Fig. S3**). Similarly, the f5C and m5C modifications are also strongly increased with aging (f5C) and tauopathy (f5C, m5C) (**Fig. 5**). Both modifications function to increase mitochondrial protein synthesis by facilitating mt-tRNA-MET initiation of translation.^93^ The changes in m1A and m1G are also potentially important. These modifications are added to 19 of 22 mt-tRNAs at position 9 by the mitochondrial ribonuclease P protein (MRPP1) Trmt10c.^94, 95^ This is likely a neuroprotective action because mutations in genes such as Trmt10c lead to neurodegenerative disorders.^9^ Thus, there is a broad-based increase in mt-tRNA level and activity that may reflect the increased turnover of mitochondria well known for aging and disease, with a resulting need for increased translation of mitochondrial proteins.^96–99^ Chronic neurodegenerative diseases, such as tauopathy, increase mitochondrial damage, which could increase the need for translation of mitochondrial proteins with a resulting requisite need for increased levels of key tRNAs.^96–99^

Perhaps the most striking link between tRNA reprogramming and codon-biased translation caused by tauopathy involve cytosolic tRNA-Arg-TCT-5-1. This isodecoder, which reads the AGA codon, increases 10-fold at 9 mo in P301S MAPT mice but not in WT mice, in parallel with increased translation of Slc, Rab, and Tmem gene families. While the Slc and Tmem families, as well as mitophagy regulator Atg9a, over-use G- and C-ending codons, the Rab family over-uses A- and T-ending codons, including AGA (**Fig. 3, Suppl. Fig. S4**). This differential codon enrichment is illustrated by a highly significant negative correlation between codons GluGAG, IleATC, LeuCTG, LysAAG, and PheTTC in Slc genes and the ArgAGA codon in Rab family genes as well as mitobiogenesis factor Nrf1 and Map2k5 (**Suppl. Fig. S4; Fig. 3**; Pearson’s p<0.05). The strongly increased level of tRNA-Arg-TCT-5-1 in neurons thus likely drives translation of critical mitobiogenesis and mitophagy genes in response to tau pathology.

It is important to note that the C57BL/6 background used for these mouse studies has a mutation in tRNA-Arg-TCT-4-1. This mutation does not impact on the mouse brain sufficiently to elicit an age-related response for Arg-TCT-5-1 in WT C57BL/6 mice, however the added stress of tauopathy disease progression is sufficient to elicit the strong response. Recent advances allow examination of tRNA isodecoders in selective brain populations.^10^ The Arg-TCT-5-1 is a isodecoder that is strongly present in neurons^10^ and focuses attention on the Arg-AGA codon. The relatively modest changes in the other tRNA isodecoders might reflect the bulk analysis used in our study and could be masking disease-related changes occurring in select cell populations.

## Materials and Methods

### Mouse Model

Use of all animals was approved by the Boston University Institutional and Animal Care and Use Committee (IACUC). All animals used in this study were handled according to IACUC approved protocols and housed in IACUC approved vivaria at the Boston University Animal Science Center. The PS19 P301S MAPT mice (B6; C3-Tg (Prnp-MAPT*P301S)PS19Vle/J, stock #008169) and C57BL/6J mice (stock #000664) were originally purchased from the Jackson Laboratory in Maine^21^ (n=4M/4F per cohort). P301S(−/−) is equivalent to wild-type (WT): no expression of human P301S tau. P301S(+/−) mice are heterozygous for human P301S mutant tau protein over-expression. Brain hemisections were obtained from euthanized littermates at 3, 6, and 9 mo of age from WT P301S(−/−) and P301S(+/−) mutant mice. Mouse and tissue information is detailed in **Supplementary Table S1**.

### RNA purification

Frozen brain tissue was transferred to a 15 mL tube and homogenized in 2 ml QiAzol using a TissueRuptor (Qiagen). A portion of this lysate (500 µL) was subjected to large (5S, 18S, 28S rRNA) and small RNA (<200 nt); miRNA, >85% tRNA) purification using the Invitrogen PureLink miRNA Isolation Kit. RNA was eluted and stored in 50 µL of RNA-free water, with concentration checked by Nanodrop (**Supplementary Table S7**). The quality of all small RNA samples was checked using an Agilent 2100 Bioanalyzer with RNA 6000 Pico chips and the results shown in **Supplementary Figure S7**.

### RNA hydrolysis

RNA from each sample (1.8 µg) was hydrolyzed in a 30 µL digestion cocktail containing 2.49 U benzonase, 3 U CIAP (calf intestinal alkaline phosphatase), 0.07 U PDE I (phosphodiesterase I), 0.1 mM deferoxamine (antioxidant), 0.1 mM butylated hydroxytoluene (antioxidant), 3 ng coformycin (adenosine deaminase inhibitor), 25 nM [^15^N]_5_-deoxyadenosine (internal standard to normalize for day-to-day variation), 2.5 mM MgCl_2_, and 5 mM Tris-HCl buffer pH 8.0. The digestion mixture was incubated at 37 °C for 6 h.

### LC-MS/MS analysis of ribonucleosides

After hydrolysis, all 54 samples were analyzed by chromatography-coupled triple-quadrupole mass spectrometry (LC-MS/MS). For each sample, 600 ng of hydrolysate was analyzed twice as two technical replicates. Using synthetic standards, HPLC retention times for the ribonucleosides were confirmed on a Waters Acuity BEH C18 column (50 × 2.1 mm inner diameter, 1.7 µm particle size) coupled to an Agilent 1290 HPLC system and an Agilent 6495 triple-quadrupole mass spectrometer. The Agilent sample vial insert was used. The HPLC system was operated at 25 °C and a flow rate of 0.3 mL/min in a gradient (**Supplementary Table S8**) with Buffer A (0.02% formic acid in water) and Buffer B (0.02% formic acid in 70% acetonitrile). The HPLC column was coupled to the mass spectrometer with an electrospray ionization source in positive mode with the following parameters: Dry gas temperature, 200 °C; gas flow, 11 L/min; nebulizer, 20 psi; sheath gas temperature, 300 °C; sheath gas flow, 12 L/min; capillary voltage, 3000 V; nozzle voltage, 0 V. As detailed in **Supplementary Table S9**, multiple reaction monitoring (MRM) mode was used for detection of product ions derived from the precursor ions for all the RNA modifications with instrument parameters including the collision energy (CE) optimized for maximal sensitivity for the modification. Signal intensities for each ribonucleoside were normalized by dividing by the sum of the UV signal intensities of the four canonical ribonucleosides recorded with an in-line UV spectrophotometer at 260 nm. For ribonucleosides lacking synthetic standards, tentative identification was based on appropriate *m/z* transitions, expected relative HPLC elution based on literature values, and, in some cases, high mass accuracy MS analysis of the eluting signal.

#### Proteomics

For protein extraction, a mixture of hippocampus and cortex were defrosted on ice, and white matter was carefully removed. Tissues were rinsed by ice-cold PBS (1x, ThermoFisher) 2 times then homogenized in lysis buffer (10mM HEPES pH8, 150 mM NaCl, 5% glycerol, 0.5% DDM, 1 mM DDT) with protease inhibitor (Sigma) and PhosSTOP (Roche) using a Dounce tissue grinder (ThermoFisher). To reduce the DNA, 2 μL of Benzonase (Sigma) was spike into the complex extract and rotated at 4°C for 30 min. The samples were centrifuged at 12,000 g for 10 min and the supernatant was transferred to a new tube. After quantification using a BCA protein assay kit (ThermoFisher), 2 mg of protein complex extract was injected to the IEX column for co-fractionation, 200 ug of protein was save for further proteomic and phosphoproteomic analysis, another 500ug of protein was kept for immunoprecipitation MS experiment. The rest of the samples were snap freezed and kept in −80°C prior to further analysis.

Purified proteins were fragmented as follows. After pelleting, the protein precipitates from the organic solvent extraction^100^ were resuspended in 250 μL of lysis buffer containing 6 M guanidine hydrochloride (GuHCl), protease inhibitors (Sigma), and phosphatase inhibitors (Roche). The samples were heated at 95°C for 10 min, cooled on ice for 10 min, and then briefly sonicated to shear the nucleic acids. The samples were diluted with 100 mM Tris (pH 8.5) to reduce the concentration of GuHCl to 0.75 M. After quantification with a BCA kit (Thermo Scientific), the proteins were digested overnight with sequence-grade trypsin (enzyme-to-protein ratio of 1:50) at 37°C, and formic acid was then added to obtain a final concentration of 1% in solution. The resulting peptides were desalted using a C18 Sep-Pak (Waters) according to the manufacturer’s instructions.

Prior to tandem mass tag (TMT) labeling, peptide quantification was performed by Pierce quantitative colorimetric assay (Thermo Scientific). According to the manufacturer’s instructions, 100 μg of peptide per sample was resuspended in 0.1 M triethylammonium bicarbonate (TEAB). Peptides of cancer cell (5 channels per condition, glucose & no glucose) were labeled with 0.2 mg of TMT 10-plex while 15 samples of human serum (5 channels per condition, healthy, pre- and post-dialysis) were labeled with TMTpro (Thermo Scientific) for 1 h at room temperature. To quench the reaction, 5% hydroxylamine was added to each sample, and the resulting mixture was incubated at room temperature for 15 min. After labeling, equal amounts of each sample were combined in a new microtube and desalted using a C18 Sep-Pak (Waters).

For proteomic data analysis, MS2 spectra were processed and searched by MaxQuant (version 1.6)^101^ against a database containing native (forward) human protein sequences (UniProt) and reversed (decoy) sequences for protein identification. The search allowed for two missed trypsin cleavage sites, variable modifications of methionine oxidation, and N-terminal acetylation. Both carbamidomethylation of cysteine residues and TMT (peptide N-termini, K residues) were set as fixed modifications. Precursor ions were searched with a maximum mass tolerance of 4.5 ppm and fragment ions with a maximum mass tolerance of 20 ppm. The candidate peptide identifications were filtered assuming a 1% FDR threshold based on searching the reverse sequence database. Quantification was performed using the TMT reporter on MS2 (TMT10-plex). Proteins with less than 30% missing value across all samples were kept for quantification. The reporter ion intensities were log-transferred and normalized using quantile normalization.

#### AQRNA-seq analysis

Absolute Quantification RNA Sequencing (AQRNA-seq) was conducted to quantify tRNA molecules at the isodecoder level in brain tissues of both P301S (−/−) and P301S (+/−) mice over the time course. cDNA libraries were constructed using 75 ng of small RNAs as input, following a revised version of the original AQRNA-seq protocol,^52, 53^ which integrates novel strategies to mitigate polymerase fall-off and primer dimers for improved quantitative performance. Specifically, the revised protocol incorporates (i) a high-processivity reverse transcriptase that is insensitive to the presence of diverse tRNA modifications during reverse transcription (RT) and (ii) a simple Exonuclease I digestion step after RT to remove excess RT primers, thereby preventing their formation from the first place. Consequently, the labor-intensive steps of AlkB demethylation and gel purification are eliminated. Compared to the original protocol, the revised version has been shown to achieve superior quantitative accuracy and sensitivity,^102^ which is particularly beneficial for detecting low-abundance isodecoders. The constructed libraries were submitted to the BioMicro Center at MIT for quality assessment using AATI Fragment Analysis (Agilent) and LightCycler 480 Real-Time PCR System (Roche), followed by sequencing on an Illumina MiSeq platform using the v3 reagent kit (Illumina) to obtain 75-bp paired-end reads. The raw sequence reads were subsequently retrieved from the server and processed using an in-house pipeline to estimate tRNA abundance at the isodecoder level.

To identify isodecoders that were significantly up- or downregulated over time in P301S(−/−) and/or P301S(+/−) mice brain tissues, differential expression analysis was performed using the DESeq2 v 1.42.0 package^103^ in R Statistical Programming Environment v 4.3.2,^104^ with raw isodecoder abundance as input. The model design included time (3, 6, or 9 mo) and genotype (P301S(−/−) or P301S +/−)) as primary variables of interest, along with their interaction term to capture genotpe-specific isodecoder abundance dynamics. In addition, the library preparation trial and sequencing flow cell were included as blocking effects to account for artifactual variability in the model outcome. Briefly, the complete workflow consists of five steps, including (i) normalization of raw abundance using the median of ratios method; (ii) estimation of dispersion parameters for each isodecoder via empirical Bayes shrinkage; (iii) fitting of a negative binomial model for each isodecoder; (iv) estimation of logarithmic fold changes using Approximate Posterior Estimation for generalized linear models;^105^ and (v) significance testing using the Wald test, with the Benjamini-Hochberg method for multiple correction.

#### Data mining, informatics, and multivariate statistics

Bioinformatic analysis of proteomics data was performed in the R statistical computing environment (version 4.3.0) using a built-in tool.^106^ GO category enrichment analysis (**Supplementary Fig. S1**) was performed using the web-based tool MetaboAnalyst 5.0 against the KEGG database (https://www.genome.jp/kegg/^107^) and the Gene Ontology database.^108^ For AQNRA-seq data, some aspects of differential expression analysis were performed using the DESeq2 v 1.42.0 package^103^ in R Statistical Programming Environment v 4.3.2.^104^

## Data Availability

Raw mass spectrometry data for proteomics and RNA modification analyses, and for AQRNA-seq analyses will be made available in public repositories. All processed data is included in supplementary tables.

## Supporting Information

- **Supplementary Table S1**: Information about mice and tissue samples. See the separate Excel spreadsheet.
- **Supplementary Table S2**: Proteomics analysis in brain hemisections from P301S(+/−) mutant mice compared to P301S(−/−) wild-type (WT) mice at 3, 6, and 9 months. See the separate Excel spreadhsheet.
- **Supplementary Table S3**: Gene-specific isoacceptor codon usage analysis for genes encoding the 100 most up- and down-regulated proteins in P301S(+/−) mice compared to wild-type P301S(−/−) mice. See the separate Excel spreadsheet.
- **Supplementary Table S4:** AQRNA-seq analysis of cytoplasmic and mitochondrial tRNAs, rRNAs, and miRNAs in brain hemisections from P301S(+/−) mutant mice compared to P301S(−/−) wild-type (WT) mice at 3, 6, and 9 months. See the separate Excel spreadsheet.
- **Supplementary Table S5**: LC-MS/MS analysis of 50 tRNA modifications in brain hemisections from P301S(+/−) mutant mice compared to P301S(−/−) wild-type (WT) mice at 3, 6, and 9 months. See the separate Excel spreadsheet.
- **Supplementary Table S6**: m1A writers and erasers
- **Supplementary Table S7**: Small RNA concentration in P301S brain tissue samples
- **Supplementary Table S8**: HPLC elution conditions for LC-MS/MS analysis
- **Supplementary Table S9:** LC-MS/MS parameters for modified ribonucleosides
- **Supplementary Figure S1**: Proteomics quality control data and identification of the 20 most up- and down-regulated proteins.
- **Supplementary Figure S2:** Analysis of age- and tauopathy-dependent changes in individual protein families.
- **Supplementary Figure S3**: Codon usage patterns in Tmem, Rab, and Slc genes up- and down-regulated by expression of mutant tau protein in the P301S mouse model.
- **Supplementary Figure S4**: Correlation analysis of codon biases in Rab and Slc genes.
- **Supplementary Figure S5**: AQRNA-seq analysis of ribosomal RNA fragments in small RNA extracted from P301S mouse brain.
- **Supplementary Figure S6:** Synthesis pathways for RNA modifications
- **Supplementary Figure S7**: Bioanalyzer tracings of small RNA isolated from P301S mouse brain tissue

## Supporting information

Sun et al Supplementary Information

Sun et al Supplementary Table S5

Sun et al Supplementary Table S4

Sun et al Supplementary Table S3

Sun et al Supplementary Table S2

Sun et al Supplementary Table S1

## References

1. Scheltens, P. et al. Alzheimer’s disease. Lancet 397, 1577–1590 (2021).

2. Romero-Molina, C., Garretti, F., Andrews, S.J., Marcora, E. & Goate, A.M. Microglial efferocytosis: Diving into the Alzheimer’s disease gene pool. Neuron 110, 3513–3533 (2022).

3. Gobom, J., Brinkmalm, A., Brinkmalm, G., Blennow, K. & Zetterberg, H. Alzheimer’s Disease Biomarker Analysis Using Targeted Mass Spectrometry. Mol Cell Proteomics 23, 100721 (2024).

4. Kumar, M. et al. Alzheimer proteopathic tau seeds are biochemically a forme fruste of mature paired helical filaments. Brain 147, 637–648 (2024).

5. Wenger, K. et al. Common mouse models of tauopathy reflect early but not late human disease. Mol Neurodegener 18, 10 (2023).

6. Wolozin, B. & Ivanov, P. Stress granules and neurodegeneration. Nat Rev Neurosci 20, 649–666 (2019).

7. Schimmel, P. The emerging complexity of the tRNA world: mammalian tRNAs beyond protein synthesis. Nat Rev Mol Cell Biol 19, 45–58 (2018).

8. Chan, P.P. & Lowe, T.M. GtRNAdb 2.0: an expanded database of transfer RNA genes identified in complete and draft genomes. Nucleic Acids Res 44, D184–189 (2016).

9. Dittmar, K.A., Goodenbour, J.M. & Pan, T. Tissue-specific differences in human transfer RNA expression. PLoS Genet 2, e221 (2006).

10. Kapur, M., Molumby, M.J., Guzman, C., Heinz, S. & Ackerman, S.L. Cell-type-specific expression of tRNAs in the brain regulates cellular homeostasis. Neuron 112, 1397–1415 e1396 (2024).

11. Suzuki, T. The expanding world of tRNA modifications and their disease relevance. Nat Rev Mol Cell Biol 22, 375–392 (2021).

12. Dedon, P.C. & Begley, T.J. Dysfunctional tRNA reprogramming and codon-biased translation in cancer. Trends Mol Med 28, 964–978 (2022).

13. Chan, C., Pham, P., Dedon, P.C. & Begley, T.J. Lifestyle modifications: coordinating the tRNA epitranscriptome with codon bias to adapt translation during stress responses. Genome Biol 19, 228 (2018).

14. Huber, S.M., Leonardi, A., Dedon, P.C. & Begley, T.J. The Versatile Roles of the tRNA Epitranscriptome during Cellular Responses to Toxic Exposures and Environmental Stress. Toxics 7 (2019).

15. Mitchener, M.M., Begley, T.J. & Dedon, P.C. Molecular Coping Mechanisms: Reprogramming tRNAs To Regulate Codon-Biased Translation of Stress Response Proteins. Acc Chem Res 56, 3504–3514 (2023).

16. Orellana, E.A., Siegal, E. & Gregory, R.I. tRNA dysregulation and disease. Nat Rev Genet 23, 651–664 (2022).

17. Lv, X., Zhang, R., Li, S. & Jin, X. tRNA Modifications and Dysregulation: Implications for Brain Diseases. Brain Sci 14 (2024).

18. Guo, W., Russo, S. & Tuorto, F. Lost in translation: How neurons cope with tRNA decoding. Bioessays 46, e2400107 (2024).

19. Blaze, J. et al. Neuronal Nsun2 deficiency produces tRNA epitranscriptomic alterations and proteomic shifts impacting synaptic signaling and behavior. Nat Commun 12, 4913 (2021).

20. Pereira, M. et al. Amyloid pathology reduces ELP3 expression and tRNA modifications leading to impaired proteostasis. Biochim Biophys Acta Mol Basis Dis 1870, 166857 (2024).

21. Yoshiyama, Y. et al. Synapse loss and microglial activation precede tangles in a P301S tauopathy mouse model. Neuron 53, 337–351 (2007).

22. Apicco, D.J. et al. Reducing the RNA binding protein TIA1 protects against tau-mediated neurodegeneration in vivo. Nat Neurosci 21, 72–80 (2018).

23. Poonaki, E., Kahlert, U.D., Meuth, S.G. & Gorji, A. The role of the ZEB1-neuroinflammation axis in CNS disorders. J Neuroinflammation 19, 275 (2022).

24. Llorens, M.C. et al. Phosphorylation Regulates Functions of ZEB1 Transcription Factor. J Cell Physiol 231, 2205–2217 (2016).

25. Li, H. et al. A dual role of human tRNA methyltransferase hTrmt13 in regulating translation and transcription. EMBO J 41, e108544 (2022).

26. Wilkinson, M.L., Crary, S.M., Jackman, J.E., Grayhack, E.J. & Phizicky, E.M. The 2’-O-methyltransferase responsible for modification of yeast tRNA at position 4. Rna 13, 404–413 (2007).

27. Marconi, G.D. et al. Epithelial-Mesenchymal Transition (EMT): The Type-2 EMT in Wound Healing, Tissue Regeneration and Organ Fibrosis. Cells 10 (2021).

28. Kaarniranta, K. et al. Autophagy in age-related macular degeneration. Autophagy 19, 388–400 (2023).

29. Yang, J. et al. Guidelines and definitions for research on epithelial-mesenchymal transition. Nat Rev Mol Cell Biol 21, 341–352 (2020).

30. Kamakura, S., Moriguchi, T. & Nishida, E. Activation of the protein kinase ERK5/BMK1 by receptor tyrosine kinases. Identification and characterization of a signaling pathway to the nucleus. J Biol Chem 274, 26563–26571 (1999).

31. Nakamura, K., Uhlik, M.T., Johnson, N.L., Hahn, K.M. & Johnson, G.L. PB1 domain-dependent signaling complex is required for extracellular signal-regulated kinase 5 activation. Mol Cell Biol 26, 2065–2079 (2006).

32. Gan-Or, Z. et al. Genetic markers of Restless Legs Syndrome in Parkinson disease. Parkinsonism Relat Disord 21, 582–585 (2015).

33. Huang, Y., Wang, P., Morales, R., Luo, Q. & Ma, J. Map2k5-Deficient Mice Manifest Phenotypes and Pathological Changes of Dopamine Deficiency in the Central Nervous System. Front Aging Neurosci 13, 651638 (2021).

34. Jo, M. et al. Inhibition of MEK5 suppresses TDP-43 toxicity via the mTOR-independent activation of the autophagy-lysosome pathway. Biochem Biophys Res Commun 513, 925–932 (2019).

35. Takayanagi, Y. & Onaka, T. Roles of Oxytocin in Stress Responses, Allostasis and Resilience. Int J Mol Sci 23 (2021).

36. Uvnas-Moberg, K., Gross, M.M., Calleja-Agius, J. & Turner, J.D. The Yin and Yang of the oxytocin and stress systems: opposites, yet interdependent and intertwined determinants of lifelong health trajectories. Front Endocrinol (Lausanne) 15, 1272270 (2024).

37. Dahlin, A., Royall, J., Hohmann, J.G. & Wang, J. Expression profiling of the solute carrier gene family in the mouse brain. J Pharmacol Exp Ther 329, 558–570 (2009).

38. Ashleigh, T., Swerdlow, R.H. & Beal, M.F. The role of mitochondrial dysfunction in Alzheimer’s disease pathogenesis. Alzheimers Dement 19, 333–342 (2023).

39. Wang, W., Zhao, F., Ma, X., Perry, G. & Zhu, X. Mitochondria dysfunction in the pathogenesis of Alzheimer’s disease: recent advances. Mol Neurodegener 15, 30 (2020).

40. Kauppila, T.E.S., Kauppila, J.H.K. & Larsson, N.G. Mammalian Mitochondria and Aging: An Update. Cell Metab 25, 57–71 (2017).

41. Popov, L.D. Mitochondrial biogenesis: An update. J Cell Mol Med 24, 4892–4899 (2020).

42. Farouk, S.M., Abdellatif, A.M. & Metwally, E. Outer and inner mitochondrial membrane proteins TOMM40 and TIMM50 are intensively concentrated and localized at Purkinje and pyramidal neurons in the New Zealand white rabbit brain. Anat Rec (Hoboken) 305, 209–221 (2022).

43. Paz, E. et al. Biochemical and neurophysiological effects of deficiency of the mitochondrial import protein TIMM50. Elife 13 (2024).

44. Tamura, H., Shibata, M., Koike, M., Sasaki, M. & Uchiyama, Y. Atg9A protein, an autophagy-related membrane protein, is localized in the neurons of mouse brains. J Histochem Cytochem 58, 443–453 (2010).

45. Markham, B.N., et al. miRNA family miR-29 inhibits PINK1-PRKN dependent mitophagy via ATG9A. bioRxiv (2024).

46. Lee, E.G., Chen, S., Leong, L., Tulloch, J. & Yu, C.E. TOMM40 RNA Transcription in Alzheimer’s Disease Brain and Its Implication in Mitochondrial Dysfunction. Genes (Basel) 12 (2021).

47. Faitg, J. et al. 3D neuronal mitochondrial morphology in axons, dendrites, and somata of the aging mouse hippocampus. Cell Rep 36, 109509 (2021).

48. Wang, X. et al. Impaired balance of mitochondrial fission and fusion in Alzheimer’s disease. J Neurosci 29, 9090–9103 (2009).

49. Li, G. & Marlin, M.C. Rab family of GTPases. Methods Mol Biol 1298, 1–15 (2015).

50. Gu, C., Begley, T.J. & Dedon, P.C. tRNA modifications regulate translation during cellular stress. FEBS Lett 588, 4287–4296 (2014).

51. Doyle, F. et al. Gene- and genome-based analysis of significant codon patterns in yeast, rat and mice genomes with the CUT Codon UTilization tool. Methods 107, 98–109 (2016).

52. Chen, R., Yim, D. & Dedon, P.C. AQRNA-seq for Quantifying Small RNAs. J Vis Exp (2024).

53. Hu, J.F. et al. Quantitative mapping of the cellular small RNA landscape with AQRNA-seq. Nat Biotechnol 39, 978–988 (2021).

54. Yu, P. et al. Dynamic Landscapes of tRNA Transcriptomes and Translatomes in Diverse Mouse Tissues. Genomics Proteomics Bioinformatics 21, 834–849 (2023).

55. Oberacker, P. et al. Bio-On-Magnetic-Beads (BOMB): Open platform for high-throughput nucleic acid extraction and manipulation. PLoS Biol 17, e3000107 (2019).

56. Natsidis, P., Schiffer, P.H., Salvador-Martinez, I. & Telford, M.J. Computational discovery of hidden breaks in 28S ribosomal RNAs across eukaryotes and consequences for RNA Integrity Numbers. Sci Rep 9, 19477 (2019).

57. Cai, W.M. et al. A Platform for Discovery and Quantification of Modified Ribonucleosides in RNA: Application to Stress-Induced Reprogramming of tRNA Modifications. Methods Enzymol 560, 29–71 (2015).

58. Hia, F. et al. Mycobacterial RNA isolation optimized for non-coding RNA: high fidelity isolation of 5S rRNA from Mycobacterium bovis BCG reveals novel post-transcriptional processing and a complete spectrum of modified ribonucleosides. Nucleic Acids Res 43, e32 (2015).

59. Li, Y. et al. 2’-O-methylation at internal sites on mRNA promotes mRNA stability. Mol Cell 84, 2320–2336 e2326 (2024).

60. Hofler, S. & Carlomagno, T. Structural and functional roles of 2’-O-ribose methylations and their enzymatic machinery across multiple classes of RNAs. Curr Opin Struct Biol 65, 42–50 (2020).

61. Zorbas, C. et al. The human 18S rRNA base methyltransferases DIMT1L and WBSCR22-TRMT112 but not rRNA modification are required for ribosome biogenesis. Mol Biol Cell 26, 2080–2095 (2015).

62. You, X.J. et al. Formation and removal of 1,N6-dimethyladenosine in mammalian transfer RNA. Nucleic Acids Res 50, 9858–9872 (2022).

63. Dominissini, D. et al. The dynamic N(1)-methyladenosine methylome in eukaryotic messenger RNA. Nature 530, 441–446 (2016).

64. Saikia, M., Fu, Y., Pavon-Eternod, M., He, C. & Pan, T. Genome-wide analysis of N1-methyl-adenosine modification in human tRNAs. RNA 16, 1317–1327 (2010).

65. Khoddami, V. & Cairns, B.R. Identification of direct targets and modified bases of RNA cytosine methyltransferases. Nat Biotechnol 31, 458–464 (2013).

66. Squires, J.E. et al. Widespread occurrence of 5-methylcytosine in human coding and non-coding RNA. Nucleic Acids Res 40, 5023–5033 (2012).

67. Enroth, C. et al. Detection of internal N7-methylguanosine (m7G) RNA modifications by mutational profiling sequencing. Nucleic Acids Res 47, e126 (2019).

68. Malbec, L. et al. Dynamic methylome of internal mRNA N(7)-methylguanosine and its regulatory role in translation. Cell Res 29, 927–941 (2019).

69. Motorin, Y., Quinternet, M., Rhalloussi, W. & Marchand, V. Constitutive and variable 2’-O-methylation (Nm) in human ribosomal RNA. RNA Biol 18, 88–97 (2021).

70. Pichot, F. et al. Quantification of substoichiometric modification reveals global tsRNA hypomodification, preferences for angiogenin-mediated tRNA cleavage, and idiosyncratic epitranscriptomes of human neuronal cell-lines. Comput Struct Biotechnol J 21, 401–417 (2023).

71. Vanderweyde, T. et al. Interaction of tau with the RNA-Binding Protein TIA1 Regulates tau Pathophysiology and Toxicity. Cell Rep 15, 1455–1466 (2016).

72. Ash, P.E.A. et al. TIA1 potentiates tau phase separation and promotes generation of toxic oligomeric tau. Proc Natl Acad Sci U S A 118 (2021).

73. Meyer, P.F. et al. Bi-directional Association of Cerebrospinal Fluid Immune Markers with Stage of Alzheimer’s Disease Pathogenesis. J Alzheimers Dis 63, 577–590 (2018).

74. Xiong, W. et al. N1-methyladenosine formation, gene regulation, biological functions, and clinical relevance. Mol Ther 31, 308–330 (2023).

75. Jin, H., Huo, C., Zhou, T. & Xie, S. m(1)A RNA Modification in Gene Expression Regulation. Genes (Basel) 13 (2022).

76. Wang, J.T. et al. Commonality and diversity in tRNA substrate recognition in t6A biogenesis by eukaryotic KEOPSs. Nucleic Acids Res 50, 2223–2239 (2022).

77. Schweizer, U., Bohleber, S. & Fradejas-Villar, N. The modified base isopentenyladenosine and its derivatives in tRNA. RNA Biology 14, 1197–1208 (2017).

78. Suzuki, T. et al. Complete chemical structures of human mitochondrial tRNAs. Nat Commun 11, 4269 (2020).

79. Auxilien, S., Guerineau, V., Szweykowska-Kulinska, Z. & Golinelli-Pimpaneau, B. The human tRNA m (5) C methyltransferase Misu is multisite-specific. RNA Biol 9, 1331–1338 (2012).

80. Tuorto, F. et al. RNA cytosine methylation by Dnmt2 and NSun2 promotes tRNA stability and protein synthesis. Nat Struct Mol Biol 19, 900–905 (2012).

81. Watanabe, Y. et al. Primary and higher order structures of nematode (Ascaris suum) mitochondrial tRNAs lacking either the T or D stem. J Biol Chem 269, 22902–22906 (1994).

82. Moriya, J. et al. A novel modified nucleoside found at the first position of the anticodon of methionine tRNA from bovine liver mitochondria. Biochemistry 33, 2234–2239 (1994).

83. Nakano, S. et al. NSUN3 methylase initiates 5-formylcytidine biogenesis in human mitochondrial tRNA(Met). Nat Chem Biol 12, 546–551 (2016).

84. Kawarada, L. et al. ALKBH1 is an RNA dioxygenase responsible for cytoplasmic and mitochondrial tRNA modifications. Nucleic Acids Res 45, 7401–7415 (2017).

85. Takemoto, C. et al. Unconventional decoding of the AUA codon as methionine by mitochondrial tRNAMet with the anticodon f5CAU as revealed with a mitochondrial in vitro translation system. Nucleic Acids Res 37, 1616–1627 (2009).

86. Suzuki, T., Nagao, A. & Suzuki, T. Human mitochondrial tRNAs: biogenesis, function, structural aspects, and diseases. Annu Rev Genet 45, 299–329 (2011).

87. Pais de Barros, J.P., et al. 2’-O-methyl-5-formylcytidine (f5Cm), a new modified nucleotide at the ‘wobble’ of two cytoplasmic tRNAs Leu (NAA) from bovine liver. Nucleic Acids Res 24, 1489–1496 (1996).

88. Feng, Y. et al. Chemical labeling - Assisted mass spectrometry analysis for sensitive detection of cytidine dual modifications in RNA of mammals. Anal Chim Acta 1098, 56–65 (2020).

89. Fu, L. et al. Tet-mediated formation of 5-hydroxymethylcytosine in RNA. J Am Chem Soc 136, 11582–11585 (2014).

90. Jiang, L. et al. Interaction of tau with HNRNPA2B1 and N(6)-methyladenosine RNA mediates the progression of tauopathy. Mol Cell 81, 4209–4227 e4212 (2021).

91. Windham, I.A. & Cohen, S. The cell biology of APOE in the brain. Trends Cell Biol 34, 338–348 (2024).

92. Ayyubova, G. Dysfunctional microglia and tau pathology in Alzheimer’s disease. Rev Neurosci 34, 443–458 (2023).

93. Lee, M. et al. Selection of initiator tRNA and start codon by mammalian mitochondrial initiation factor 3 in leaderless mRNA translation. Nucleic Acids Res 53 (2025).

94. Vilardo, E. et al. A subcomplex of human mitochondrial RNase P is a bifunctional methyltransferase--extensive moonlighting in mitochondrial tRNA biogenesis. Nucleic Acids Res 40, 11583–11593 (2012).

95. Meynier, V. et al. Structural basis for human mitochondrial tRNA maturation. Nat Commun 15, 4683 (2024).

96. Yang, H.M. Mitochondrial Dysfunction in Neurodegenerative Diseases. Cells 14 (2025).

97. Antico, O., Thompson, P.W., Hertz, N.T., Muqit, M.M.K. & Parton, L.E. Targeting mitophagy in neurodegenerative diseases. Nat Rev Drug Discov 24, 276–299 (2025).

98. Rummel, N.G. & Butterfield, D.A. Altered Metabolism in Alzheimer Disease Brain: Role of Oxidative Stress. Antioxid Redox Signal 36, 1289–1305 (2022).

99. Zhu, X., Smith, M.A., Perry, G. & Aliev, G. Mitochondrial failures in Alzheimer’s disease. Am J Alzheimers Dis Other Demen 19, 345–352 (2004).

100. Causon, T.J. & Hann, S. Review of sample preparation strategies for MS-based metabolomic studies in industrial biotechnology. Anal Chim Acta 938, 18–32 (2016).

101. Tyanova, S., Temu, T. & Cox, J. The MaxQuant computational platform for mass spectrometry-based shotgun proteomics. Nat Protoc 11, 2301–2319 (2016).

102. Chen, R., Yim, D., Liu, L., Cao, B. & Dedon, P. Toward automated NGS RNA-seq for absolute quantification of small RNAs: A side-by-side comparison of library preparation protocols. bioRxiv (2024).

103. Love, M.I., Huber, W. & Anders, S. Moderated estimation of fold change and dispersion for RNA-seq data with DESeq2. Genome Biol 15, 550 (2014).

104. Team, R.C. (R Foundation for Statistical Computing, Vienna, Austria; 2013).

105. Zhu, A., Ibrahim, J.G. & Love, M.I. Heavy-tailed prior distributions for sequence count data: removing the noise and preserving large differences. Bioinformatics 35, 2084–2092 (2019).

106. Blum, B.C. & Emili, A. Omics Notebook: robust, reproducible and flexible automated multiomics exploratory analysis and reporting. Bioinform Adv 1, vbab024 (2021).

107. Kanehisa, M., Sato, Y., Kawashima, M., Furumichi, M. & Tanabe, M. KEGG as a reference resource for gene and protein annotation. Nucleic Acids Res 44, D457–462 (2016).

108. Gene Ontology, C. The Gene Ontology resource: enriching a GOld mine. Nucleic Acids Res 49, D325–D334 (2021).

