## Supplementary material for "Distinct age- and pathology-dependent epitranscriptome and translational dysfunction in a tauopathy mouse model": Sun et al Supplementary Information

***Contents***


- **Supplementary Table S1**: Information about mice and tissue samples. See the separate Excel spreadsheet.
- **Supplementary Table S2**: Proteomics analysis in brain hemisections from P301S(+/-) mutant mice compared to P301S(-/-) wild-type (WT) mice at 3, 6, and 9 months. See the separate Excel spreadhsheet.
- **Supplementary Table S3**: Gene-specific isoacceptor codon usage analysis for genes encoding the 100 most up- and down-regulated proteins in P301S(+/-) mice compared to wild-type P301S(-/-) mice. See the separate Excel spreadsheet.
- **Supplementary Table S4:** AQRNA-seq analysis of cytoplasmic and mitochondrial tRNAs, rRNAs, and miRNAs in brain hemisections from P301S(+/-) mutant mice compared to P301S(-/-) wild-type (WT) mice at 3, 6, and 9 months. See the separate Excel spreadsheet.
- **Supplementary Table S5**: LC-MS/MS analysis of 50 tRNA modifications in brain hemisections from P301S(+/-) mutant mice compared to P301S(-/-) wild-type (WT) mice at 3, 6, and 9 months. See the separate Excel spreadsheet.
- **Supplementary Table S6**: m1A writers and erasers
- **Supplementary Table S7**: Small RNA concentration in P301S brain tissue samples
- **Supplementary Table S8**: HPLC elution conditions for LC-MS/MS analysis
- **Supplementary Table S9:** LC-MS/MS parameters for modified ribonucleosides

- **Supplementary Figure S1**: Proteomics quality control data and identification of the 20 most up- and down-regulated proteins.
- **Supplementary Figure S2:** Analysis of age- and tauopathy-dependent changes in individual protein families.
- **Supplementary Figure S3**: Codon usage patterns in Tmem, Rab, and Slc genes up- and down-regulated by expression of mutant tau protein in the P301S mouse model.
- **Supplementary Figure S4**: Correlation analysis of codon biases in Rab and Slc genes.
- **Supplementary Figure S5**: AQRNA-seq analysis of ribosomal RNA fragments in small RNA extracted from P301S mouse brain.
- **Supplementary Figure S6:** Synthesis pathways for RNA modifications
- **Supplementary Figure S7**: Bioanalyzer tracings of small RNA isolated from P301S mouse brain tissue

**Supplementary Table S6:** m1A writers and erasers

| ***Writers*** | **Substrate** | **Change in enzyme level** | **Refs** |
|---|---|---|---|
| Trmt10c | m1A58 in mt-tRNAs | Increased in mutant vs WT at 3, 6, 9 mo | **1** |
| TRMT6/TRMT61A | m1A58 in cy-tRNAs | Not detected | **1** |
| Bmt2 | 28S rRNA m1A3,301 in mouse | Not detected | **1** |
| Trmt61B | 16S mt-rRNA m1A947, m1A58 mt-tRNAs | Not detected | **1** |
| ***Erasers*** | | | |
| ALKBH1 | m1A58 in specific cytosolic, mitochondrial tRNAs | Not detected | **1, 2** |
| ALKBH3 | mRNA, specific tRNAs | Increase in mutant vs WT at 9 mo; in WT at 9 mo | **1** |
| ALKBH7 | | Decreased in mutant vs WT at 6 mo | **1** |
| FTO | mRNA | Decrease at 3 mo | **1** |

#### References


1. Xiong, W. et al. N1-methyladenosine formation, gene regulation, biological functions, and clinical relevance. *Mol Ther* **31**, 308-330 (2023).
2. Kawarada, L. et al. ALKBH1 is an RNA dioxygenase responsible for cytoplasmic and mitochondrial tRNA modifications. *Nucleic Acids Res* **45**, 7401-7415 (2017).

**Supplementary Table S7:** Small RNA (<200 nt) was isolated from brain hemisections from P301S (+/-) and (-/-) mice. Note: 12-13 samples were available for 3 mo and 9 mo mice, but only 2 samples for the 9-10 mo mice. Sample numbering from **Supplementary Table S1.**

| Sample | Age | Strain | RNA concentration (ng/µL) |
|---|---|---|---|
| 61 | 3 months | P301S -/- | 149.4 |
| 64 | | P301S -/- | 194.1 |
| 65 | | P301S -/- | 197.3 |
| 69 | | P301S -/- | 179.1 |
| 70 | | P301S -/- | 179.2 |
| 71 | | P301S -/- | 190.7 |
| 72 | | P301S -/- | 149.5 |
| 73 | | P301S -/- | 189.4 |
| 74 | | P301S -/- | 159 |
| 75 | | P301S -/- | 157.5 |
| 76 | | P301S -/- | 121.6 |
| 77 | | P301S -/- | 124.7 |
| 78 | | P301S -/- | 129.4 |
| 54 | 3 months | P301S +/- | 216.1 |
| 55 | | P301S +/- | 166 |
| 56 | | P301S +/- | 155.4 |
| 57 | | P301S +/- | 160.7 |
| 58 | | P301S +/- | 175.3 |
| 59 | | P301S +/- | 172.9 |
| 60 | | P301S +/- | 91.8 |
| 62 | | P301S +/- | 187.4 |
| 63 | | P301S +/- | 167 |
| 66 | | P301S +/- | 156 |
| 67 | | P301S +/- | 164.2 |
| 68 | | P301S +/- | 187.7 |
| 24 | 6 months | P301S -/- | 147.9 |
| 25 | | P301S -/- | 205.6 |
| 26 | | P301S -/- | 163 |
| 27 | | P301S -/- | 189.7 |
| 28 | | P301S -/- | 164.3 |
| 31 | | P301S -/- | 151.8 |
| 33 | | P301S -/- | 169.6 |
| 34 | | P301S -/- | 192.2 |
| 35 | | P301S -/- | 171.1 |
| 36 | | P301S -/- | 155.2 |
| 41 | | P301S -/- | 166.2 |
| 46 | | P301S -/- | 175.1 |
| 29 | 6 months | P301S +/- | 168.9 |
| 30 | | P301S +/- | 174.3 |
| 32 | | P301S +/- | 187.6 |
| 37 | | P301S +/- | 195.2 |
| 38 | | P301S +/- | 174.6 |
| 39 | | P301S +/- | 152.4 |
| 40 | | P301S +/- | 180.8 |
| 42 | | P301S +/- | 178.5 |
| 43 | | P301S +/- | 181.7 |
| 44 | | P301S +/- | 187 |
| 45 | | P301S +/- | 186.5 |
| 22 | | P301S +/- | 216 |
| 23 | | P301S +/- | 192.9 |
| 20 | 9-10 months | P301S -/- | 195.1 |
| 21 | | P301S -/- | 67.7 |
| 14 | 9-10 months | P301S +/- | 86 |
| 18 | | P301S +/- | 93.3 |

**Supplementary Table S8.** The buffer gradient used in the LC-MS/MS. Buffer A: 0.02% formic acid in double distilled water. Buffer B: 0.02% formic acid in 70% acetonitrile.

| Time (min) | Buffer A (%) | Buffer B (%) | Flow (ml/min) | Time (min) | Buffer A (%) | Buffer B (%) | Flow (ml/min) |
|---|---|---|---|---|---|---|---|
| 0 | 100 | 0 | 0.3 | 13 | 85 | 15 | 0.3 |
| 5 | 99 | 1 | 0.3 | 15 | 80 | 20 | 0.3 |
| 6 | 98 | 2 | 0.3 | 16 | 25 | 75 | 0.3 |
| 7 | 97 | 3 | 0.3 | 17 | 0 | 100 | 0.3 |
| 8 | 95 | 5 | 0.3 | 18 | 0 | 100 | 0.3 |
| 9 | 93 | 7 | 0.3 | 20 | 0 | 100 | 0.3 |
| 10 | 90 | 10 | 0.3 | 21 | 100 | 0 | 0.3 |
| 12 | 88 | 12 | 0.3 | 25 | 100 | 0 | 0.3 |

**Supplementary Table S9.** List of modified ribonucleosides included in the LC-MS/MS analysis method. Signals of several modifications were confirmed by both qualifier and quantifier transitions. Abbreviations according to http://genesilico.pl/modomics/modifications. Modifications in red font are tentative identifications that lack synthetic standards but have LC-MS/MS precedent in the literature.{Huber, 2022 #8100;Suzuki, 2021 #7998;Chujo, 2021 #9223}

| Name | Precursor Ion | Product Ion | RT (min) | Name | Precursor Ion | Product Ion | RT (min) |
|---|---|---|---|---|---|---|---|
| 15N-dA | 257 | 141 | 4.47 | m$^{5}$U | 259 | 127 | 4.10 |
| ac$^{4}$C | 286 | 154 | 7.80 | m$^{5}$Um | 273 | 127 | 9.00 |
| acp$^{3}$U | 346 | 214 | 1.36 | m$^{6,6}$A | 296 | 164 | 11.35 |
| Am | 282 | 136 | 7.32 | m$^{6}$A | 282 | 150 | 8.84 |
| Cm | 258 | 112 | 2.58 | m$^{6}$t$^{6}$A | 427 | 295 | 14.29 |
| cm$^{5}$s$^{2}$U | 319 | 187 | 6.89 | m$^{7}$G | 298 | 166 | 2.15 |
| cm$^{5}$U | 303.1 | 171 | 2.97 | man-Q | 572 | 163 | 5.48 |
| D | 247 | 115 | 0.88 | gal-Q | 572 | 295 | 6.44 |
| f$^{5}$C | 272.1 | 140.1 | 4.18 | mchm$^{5}$U *R* | 333.1 | 183 | 4.17 |
| f$^{5}$Cm | 286.1 | 140.1 | 9.30 | mchm$^{5}$U *S* | 333.1 | 183 | 6.17 |
| Gm | 298 | 152 | 7.10 | mcm$^{5}$s$^{2}$U | 333 | 201 | 10.41 |
| hm$^{5}$C | 274.1 | 142.1 | 0.90 | mcm$^{5}$U | 317.2 | 185.1 | 7.70 |
| hm$^{5}$Cm | 288 | 142.1 | 2.82 | mcm$^{5}$Um | 331.1 | 185.1 | 10.61 |
| ho$^{5}$U | 261 | 129 | 1.17 | ms$^{2}$i$^{6}$A | 382.2 | 250.1 | 17.08 |
| I | 269 | 137 | 3.86 | ms$^{2}$t$^{6}$A | 459 | 327 | 14.78 |
| i$^{6}$A | 336 | 204 | 16.49 | ncm$^{5}$U | 302 | 170 | 1.41 |
| m$^{1}$A | 282 | 150 | 1.32 | ncm$^{5}$Um | 316 | 170 | 4.10 |
| m$^{1}$G | 298 | 166 | 7.13 | oHyW | 525 | 393 | 14.42 |
| m$^{1}$I | 283 | 151 | 7.34 | Q | 410 | 163 | 5.27 |
| m$^{2,2}$G | 312 | 180 | 9.66 | s$^{2}$U | 261 | 129 | 4.36 |
| m$^{2}$G | 298 | 166 | 7.92 | t$^{6}$A | 413 | 281 | 12.50 |
| m$^{3}$C | 258 | 126 | 1.15 | Um | 259 | 113 | 5.61 |
| m$^{5}$C | 258 | 126 | 1.36 | Y | 245 | 191 | 0.89 |

**Supplementary Figure S1**: Proteomics quality control data and identification of the 20 most up- and down-regulated proteins at each age of the mutant P301S(+/-) mice compared to wild-type P301S(-/-) mice. (**A**) Gene Ontology (GO) categories represented in the proteomic dataset in **Supplementary Table S2**. (**B**) Number of up- and down-regulated proteins at p<0.05 and p<0.1 based on a Student's T-test. (**C**) Identities of the 20 most up-and down-regulated proteins at p<0.1 from **Supplementary Table S2**.

**A**

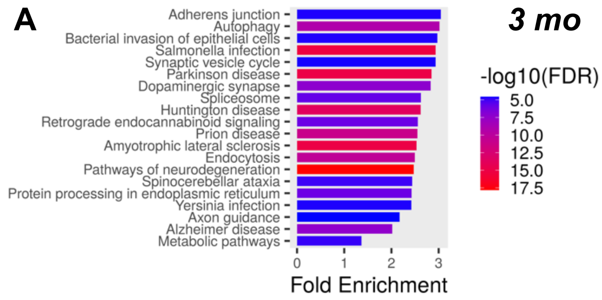


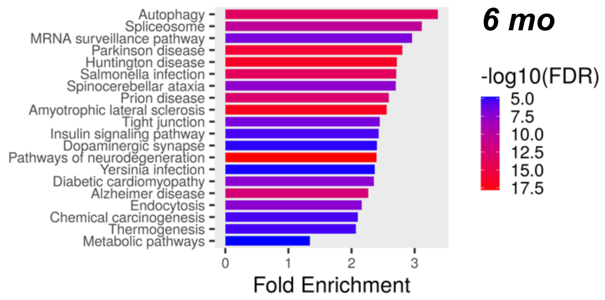


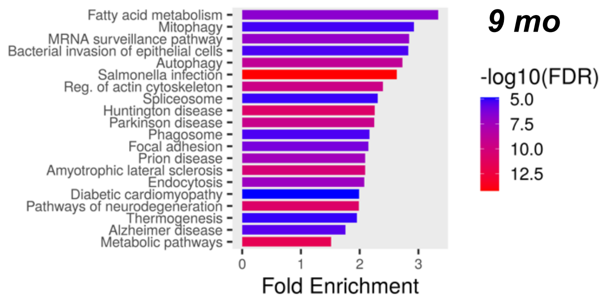


**B**

**Number of up- and down-regulated proteins in P301S(+/-) compared to P301S(-/-) mice**

| | p < 0.05 | | | p < 0.1 | | |
|---|---|---|---|---|---|---|
| Age | Up | Down | Sum | Up | Down | Sum |
| 3 mo | 20 | 16 | 36 | 1205 | 1120 | 2325 |
| 6 mo | 36 | 27 | 63 | 1264 | 1256 | 2520 |
| 9 mo | 66 | 24 | 90 | 1324 | 1535 | 2859 |

**C**

***20 Most Up- and Down-regulated Proteins at p<0.1***
***P301S (+/-) vs (-/-)***

| Top 20 Up-regulated | | | | | | Top 20 Down-regulated | | | | | |
|---|---|---|---|---|---|---|---|---|---|---|---|
| Gene names | 3 mo_log2FC | Gene names | 6 mo_log2FC | Gene names | 9 mo_log2FC | Gene names | 3 mo_log2FC | Gene names | 6 mo_log2FC | Gene names | 9 mo_log2FC |
| Ptdss2 | 3.675 | Ptdss2 | 3.457 | Ptdss2 | 3.418 | Slc1a4 | -0.473 | Med24 | -0.524 | Fbxo30 | -0.494 |
| MAPT | 1.873 | Oxt | 2.156 | C1qa | 1.753 | Fem1b | -0.475 | Lcorl | -0.524 | Acp1 | -0.499 |
| Oxt | 1.200 | MAPT | 1.918 | MAPT | 1.677 | Nudt18 | -0.485 | Fbxo25 | -0.529 | Slu7 | -0.504 |
| Rilpl2 | 0.903 | Map2k5 | 1.212 | Lag3 | 1.426 | Slc38a3 | -0.486 | Rab39b | -0.538 | Mpped2 | -0.507 |
| Map2k5 | 0.752 | Stard3 | 1.110 | C1qc | 1.311 | Cldn10 | -0.486 | Col6a2 | -0.549 | Jarid2 | -0.508 |
| Trmt13 | 0.669 | Med12 | 0.859 | Ifit3 | 1.292 | Cldn11 | -0.494 | Snrpb | -0.560 | Btf3l4 | -0.509 |
| Med12 | 0.658 | Rilpl2 | 0.853 | Adra2a | 1.255 | Rab39b | -0.514 | Atg9a | -0.562 | Clic6 | -0.514 |
| Rbm19 | 0.611 | Pdgfc | 0.759 | Cd14 | 1.171 | S100a11 | -0.514 | Hpcal4 | -0.577 | Npc1 | -0.520 |
| Myh7b | 0.593 | Rpl37 | 0.754 | Sec61g | 1.166 | Rab3d | -0.516 | Slc1a3 | -0.587 | Rapgefl1 | -0.540 |
| Rpl37 | 0.564 | Smim19 | 0.696 | C1qb | 1.157 | Slc6a3 | -0.516 | Rer1 | -0.590 | Prr3 | -0.543 |
| Msrb3 | 0.554 | Hcn3 | 0.679 | Gfap | 1.085 | Vcpkmt | -0.520 | Cldn10 | -0.594 | Acat2 | -0.553 |
| Pttg1 | 0.543 | Trmt13 | 0.677 | Lyz2 | 0.943 | Atg9a | -0.531 | Fars2 | -0.618 | Pttg1 | -0.555 |
| Trh | 0.536 | Pttg1 | 0.663 | Ifitm3 | 0.939 | Ocln | -0.531 | Ipo11 | -0.654 | Nudt17 | -0.562 |
| Celf6 | 0.532 | Msrb3 | 0.648 | Rilpl2 | 0.917 | Maf1 | -0.541 | Shprh | -0.663 | Fat4 | -0.563 |
| Eya4 | 0.526 | Myh7b | 0.627 | S100a4 | 0.907 | Vopp1 | -0.541 | Slc6a3 | -0.676 | Camta1 | -0.588 |
| Rpl34 | 0.513 | Dusp28 | 0.613 | Bst2 | 0.875 | Slc2a1 | -0.632 | Ergic3 | -0.749 | Trh | -0.600 |
| Pdgfc | 0.510 | Eya4 | 0.601 | Vcpkmt | 0.873 | Hsdl1 | -0.667 | Gtf2e2 | -0.778 | Atp11c | -0.615 |
| Dpysl3 | 0.504 | Cck | 0.600 | Pgap1 | 0.853 | Slc1a3 | -0.669 | Slc8a3 | -0.822 | Cep350 | -0.651 |
| Camkmt | 0.497 | Akt1s1 | 0.598 | C4b | 0.842 | Bcan | -0.820 | Maf1 | -0.837 | Rpl37 | -0.680 |
| Sp8 | 0.485 | Tmcc3 | 0.597 | Rasa4 | 0.833 | Fam160b2 | -0.897 | Btbd17 | -0.842 | Pdgfc | -0.703 |

**Supplementary Figure S2:** Analysis of age- and tauopathy-dependent changes in individual protein families. **(A)** Changes in the numbers of members of the Rab, Slc, and ribosomal protein families in mice expression the mutant tau protein (Mutant) as a function of age. **(B)** Quantities of cytosolic large (Rpl) and small (Rps) ribosomal proteins and mitochondrial large (Mrpl) and small (Mrps) ribosomal proteins in wild-type P301S(-/-) (WT) and mutant tau-expressing P301S(+/-) mice (Mut). Data from **Suppl. Table S2.**

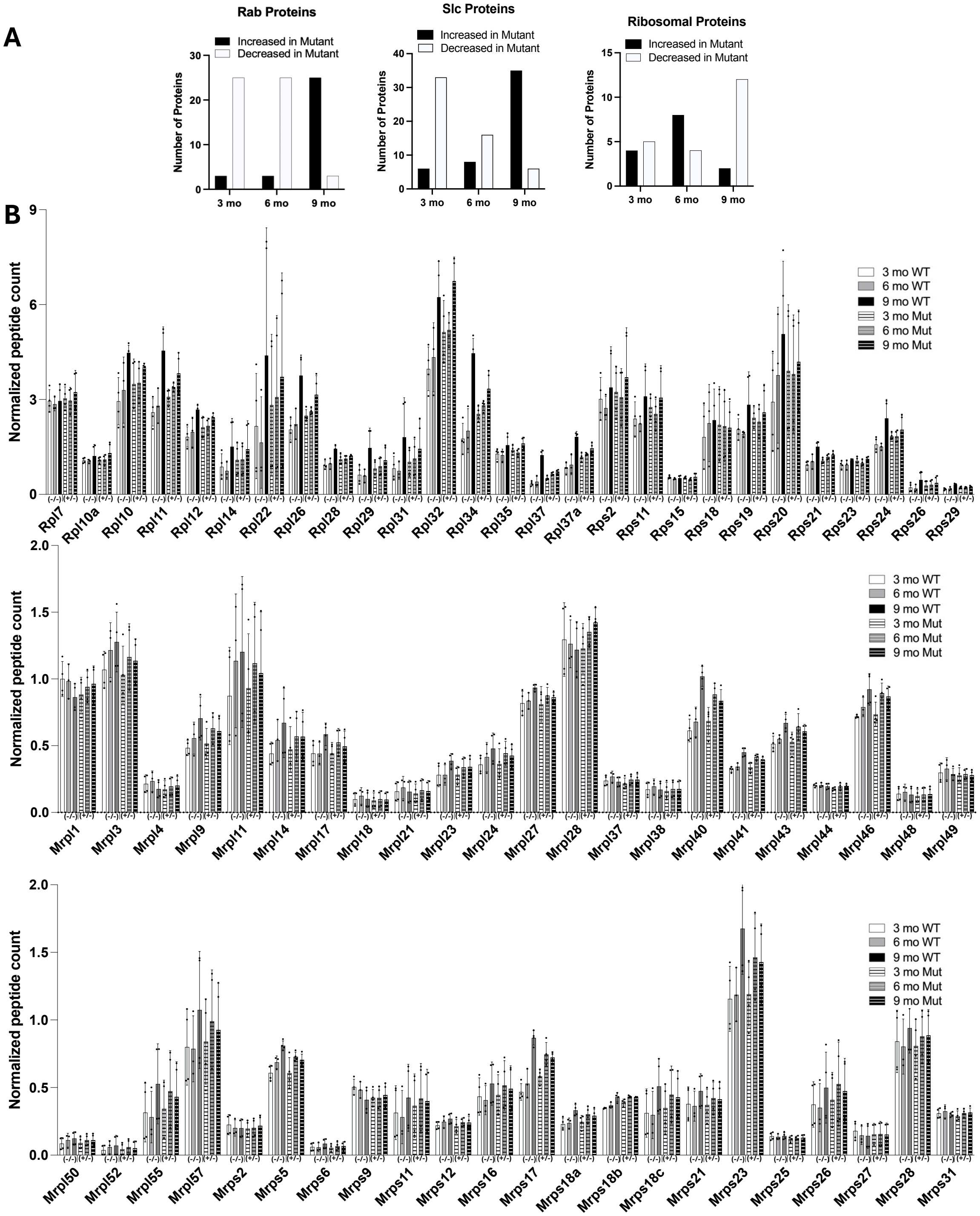

**Supplementary Figure S3**: Codon usage patterns in Tmem, Rab, and Slc genes up- and down-regulated by over-expression of mutant tau protein in the P301S mouse model. Hierarchical clustering of Z-scores for codon usage (**Suppl. Table S3**) for examples of genes up- and down-regulated in mutant P301S(+/-) relative to wild-type P301S(-/-) mice.

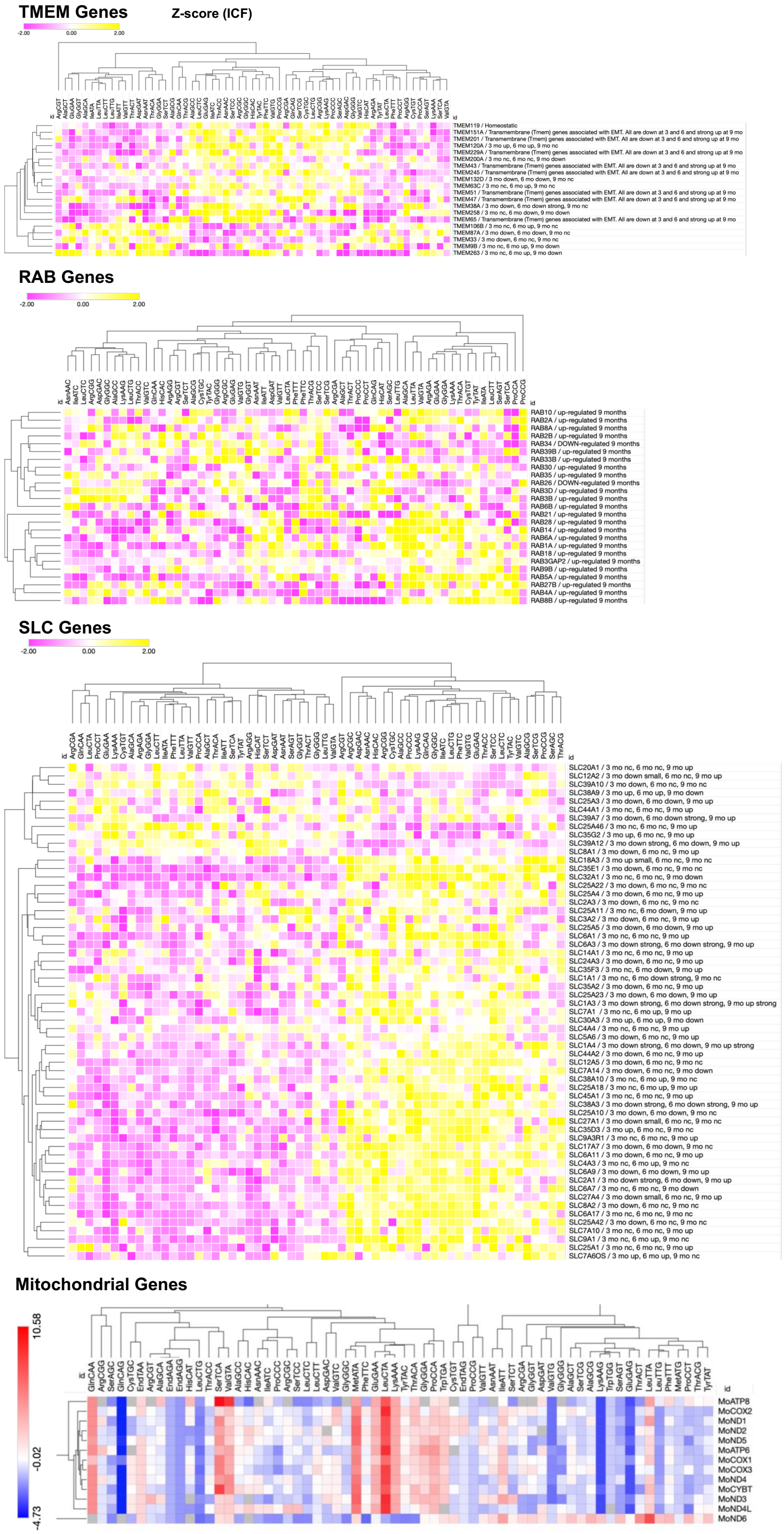

**Supplementary Figure S4:** Correlation analysis for codon biases in Rab and Slc family genes.

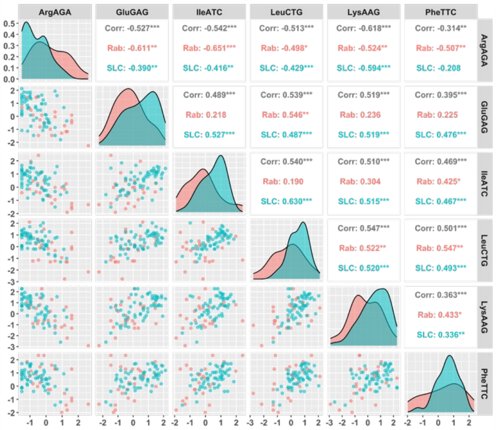


**Correlogram displaying pairwise relationships between codon usage biases of select codons in Rab and SLC genes.** The upper right triangle shows Spearman's correlation coefficients for both sets of genes (black), Rab genes (red), and SLC genes (blue). Asterisks denote significance (p < 0.05 *, p < 0.01 **, p < 0.001 ***). On the diagonal are density plots for each codon for both sets of genes, colored as above. The lower left triangle shows scatterplots of Z-scores for each codon compared to the one on the intersecting axis.

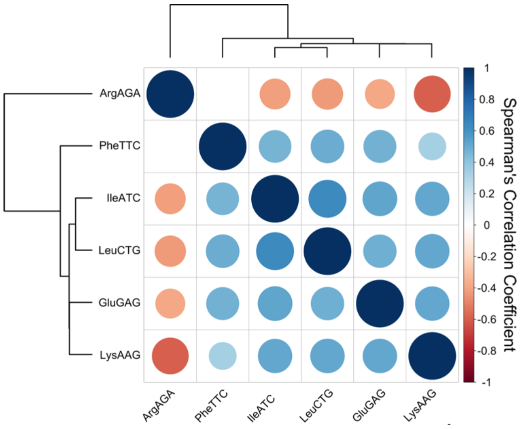


**Spearman's correlation matrix of codon usage bias for SLC genes.** Spearman's correlations coefficients were calculated between Z-scores for selected codons for all SLC genes. The size and intensity of color shows the strength of the correlation. Dots are gated by BH-adjusted p-values (adjusted p < 0.05), and only significant correlations are shown. Unsupervised hierarchical clustering was performed on all codons, represented by the dendrograms at the top and left of the image.

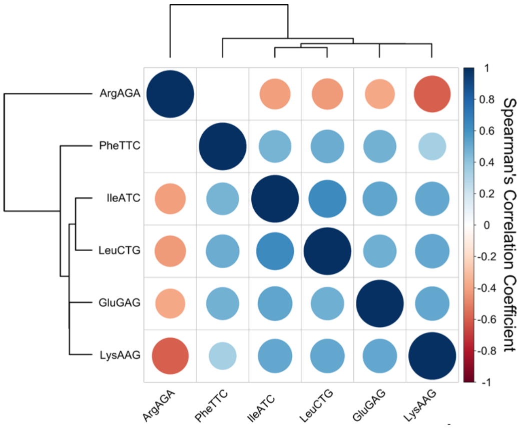


**Spearman's correlation matrix of codon usage bias for SLC genes.** Spearman's correlations coefficients were calculated between Z-scores for selected codons for all SLC genes. The size and intensity of color shows the strength of the correlation. Dots are gated by BH-adjusted p-values (adjusted p < 0.05), and only significant correlations are shown. Unsupervised hierarchical clustering was performed on all codons, represented by the dendrograms at the top and left of the image.

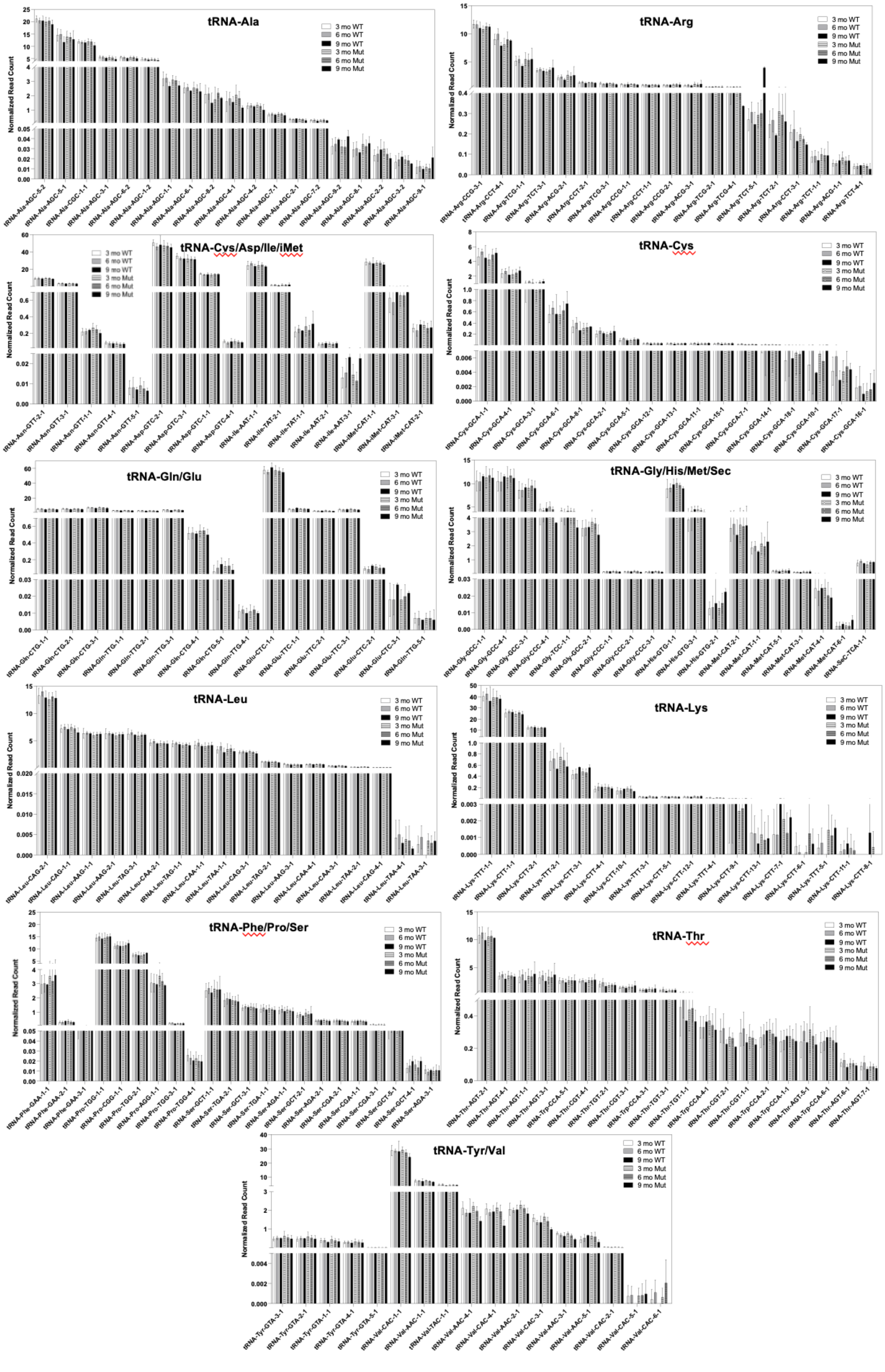
tRNA-Ala
tRNA-Arg
tRNA-Cys/Asp/Ile/iMet
tRNA-Cys
tRNA-Gln/Glu
tRNA-Gly/His/Met/Sec
tRNA-Leu
tRNA-Lys
tRNA-Phe/Pro/Ser
tRNA-Thr
tRNA-Tyr/Val
3 mo WT
6 mo WT
9 mo WT
3 mo Mut
6 mo Mut
9 mo Mut
Normalized Read Count

**Above -- Supplementary Figure S5:** AQRNA-seq analysis of tRNA isodecoders in small RNA extracted from P301S mouse brain hemisections. Graphs show the normalized read counts obtained by dividing raw reads by total RNA reads in each sample (Suppl. Table S4). Data represent mean ±SD for N=12-13 for the 3 mo and 6 mo mice and mean ±deviation about the mean for the 9 mo mice (N=2).

**Supplementary Figure S6:** Synthesis pathways for RNA modifications.

### Wobble U modifications by Elps, Cut1/2, Alkbh8, Ftsj1

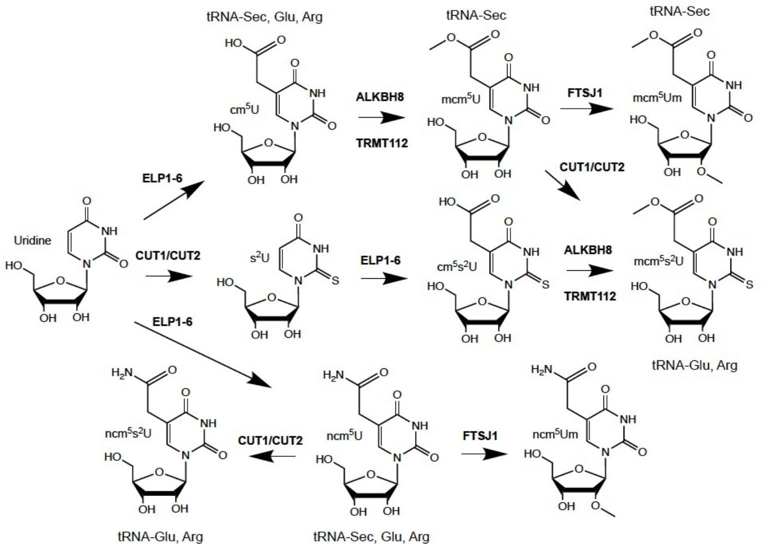

**Supplementary Figure S7:** Agilent Bioanalyzer tracings using RNA 6000 Pico chips for the 54 small RNA samples from **Supplementary Table S1** (same sample numbering with m-XX).

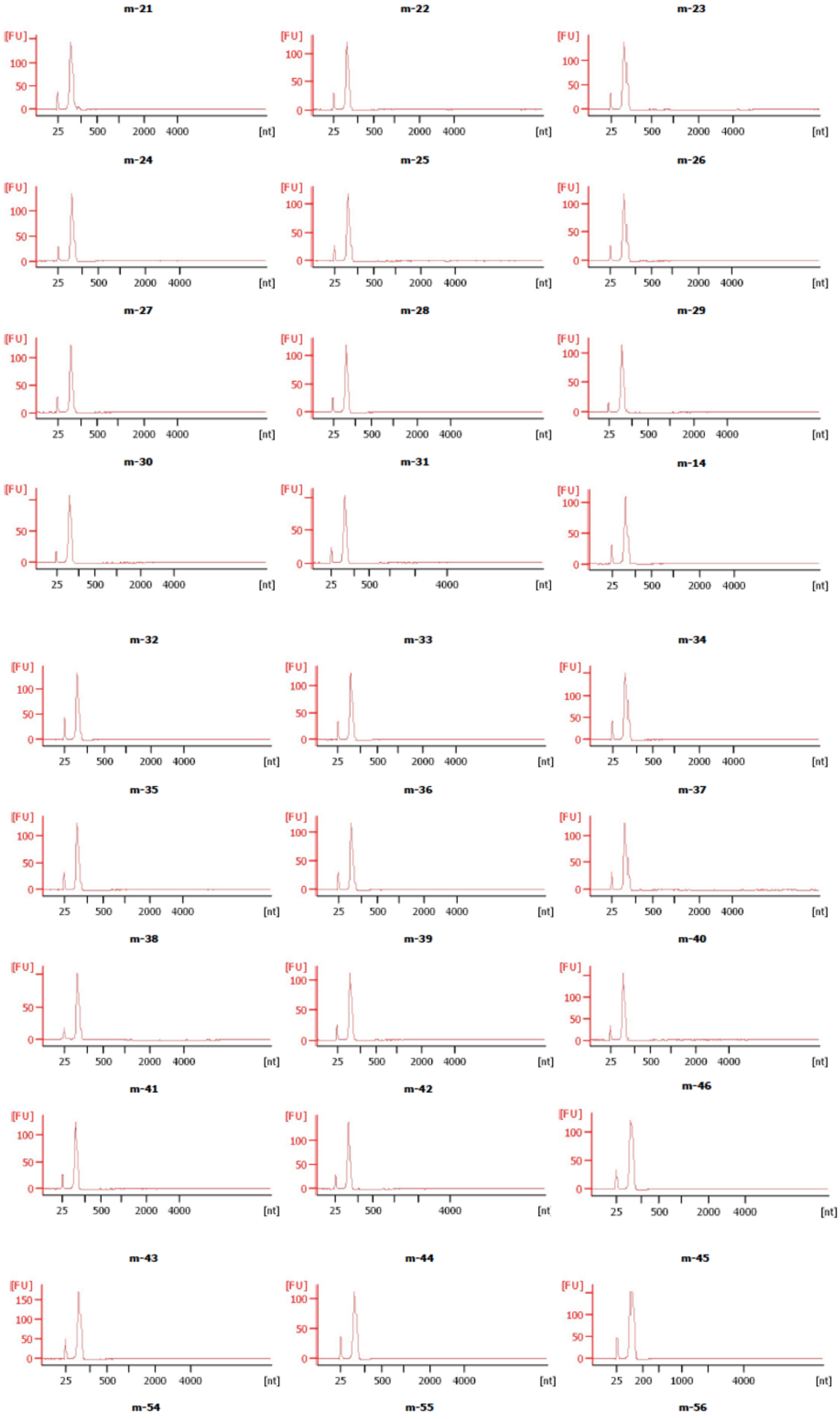

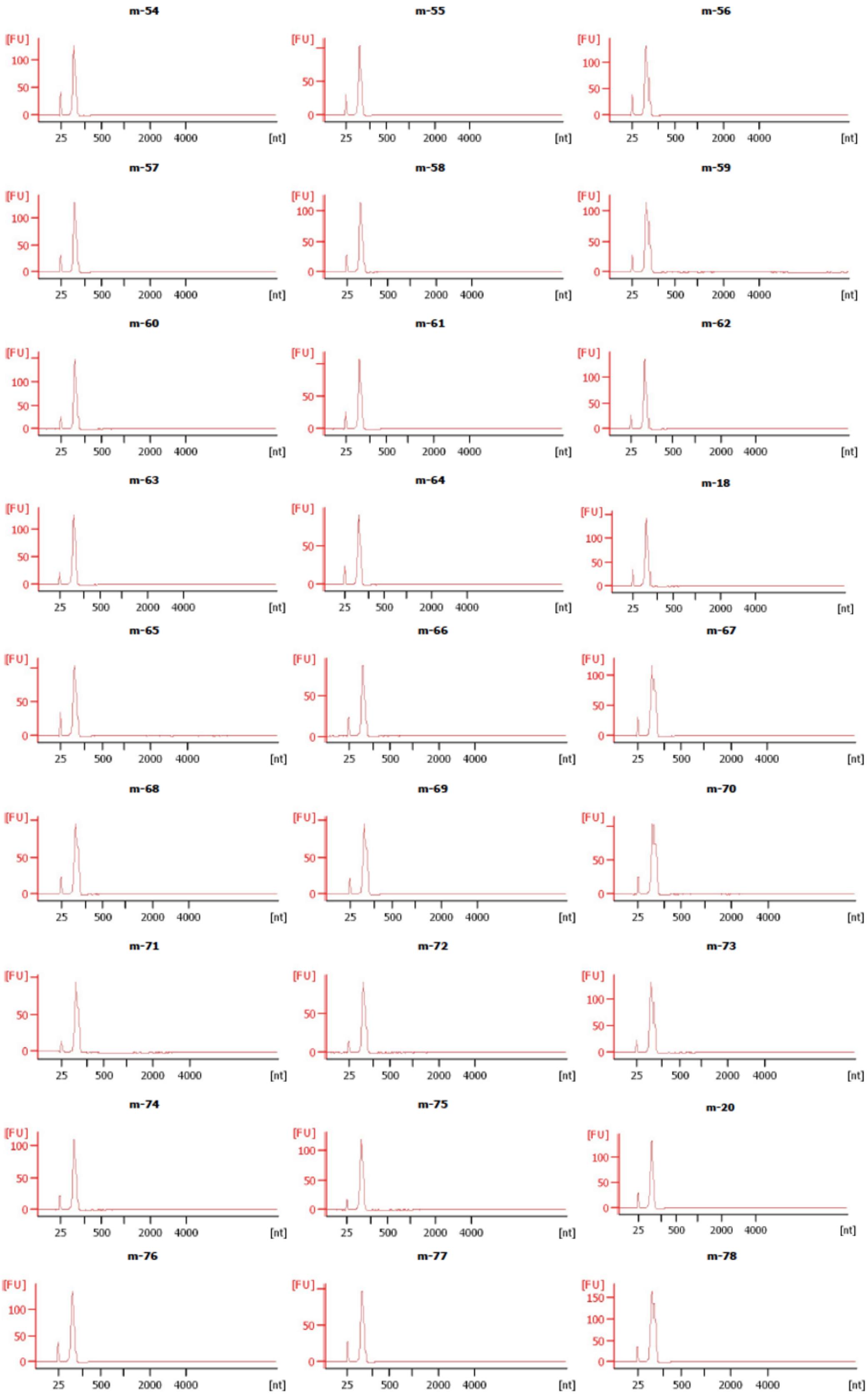
m-54
m-55
m-56
m-57
m-58
m-59
m-60
m-61
m-62
m-63
m-64
m-18
m-65
m-66
m-67
m-68
m-69
m-70
m-71
m-72
m-73
m-74
m-75
m-20
m-76
m-77
m-78
[FU]
[nt]
25
500
2000
4000
